# Resolving Between Novelty and Homology in the Rapidly Evolving Phallus of Drosophila

**DOI:** 10.1101/2021.04.14.439817

**Authors:** Gavin R. Rice, Jean R. David, Nicolas Gompel, Amir Yassin, Mark Rebeiz

## Abstract

The genitalia present some of the most rapidly evolving anatomical structures in the animal kingdom, possessing a variety of parts that can distinguish recently diverged species. In the *Drosophila melanogaster* group, the phallus is adorned with several processes, pointed outgrowths, that are similar in size and shape between species. However, the complex three-dimensional nature of the phallus can obscure the exact connection points of each process. Previous descriptions based upon adult morphology have primarily assigned phallic processes by their approximate positions in the phallus and have remained largely agnostic regarding their homology relationships. In the absence of clearly identified homology, it can be challenging to model when each structure first evolved. Here, we employ a comparative developmental analysis of these processes in eight members of the *melanogaster* species group to precisely identify the tissue from which each process forms. Our results indicate that adult phallic processes arise from three pupal primordia in all species. We found that in some cases the same primordia generate homologous structures whereas in other cases, different primordia produce phenotypically similar but remarkably non-homologous structures. This suggests that the same gene regulatory network may have been redeployed to different primordia to induce phenotypically similar traits. Our results highlight how traits diversify and can be redeployed, even at short evolutionary scales.

**Research Highlight:** By incorporating developmental analysis, we find that genital structures previously identified as homologs are novel structures. This highlights how developmental analysis can help resolve contentious claims of homology.

## Introduction

Most studies of developmental evolution depend on our ability to precisely compare the same body parts in different species or populations. Rigorously establishing such homology relationships allows us to identify novel traits, which are often defined by their lack of homology (Reviewed in Moczek, 2008; G. Wagner, 2007). Many of the current model systems for the study of novelty focus on traits that arose in the distant past (Bruce & Patel, 2020; Clark-Hachtel & Tomoyasu, 2020; Emlen, Szafran, Corley, & Dworkin, 2006; Hinman, Nguyen, Cameron, & Davidson, 2003), making the investigation of their molecular origins difficult. These traits likely arose through a multitude of genetic changes and exist in organisms that are less amenable to genetic studies. Recently evolved traits found in the rapidly evolving tissues of model organisms can provide qualitative changes in morphology produced by genomes that are easily compared and modified. However, rapidly evolving anatomical structures pose a distinct challenge. When differences between the anatomy of two species are numerous, it can be difficult to disentangle which structures are ancestral and which represent newly derived novelties. Thus, while macroevolutionary novelties often appear as clear discontinuities in the evolutionary record, the more molecularly tractable micro- and mesoevolutionary novelties require us to consider their relationships in a rich and complicated comparative context (Abouheif, 2008; Church & Extavour, 2020). Overcoming this challenge is thus critical to develop a genetic portrait of morphological novelty.

Most assertions of homology are defined through establishing that the structure in question connects to an unambiguously homologous tissue in both species (Moczek, 2008). Contentious claims of homology often revolve around the question of whether a set of traits are formed by the same cells or tissues. These assertions can be strengthened through developmental analysis, where the primordium that initially forms the trait in question can be determined (Tanaka, Barmina, & Kopp, 2009). This type of analysis is especially important in complex three-dimensional traits, as resolution in the X, Y, and Z axes may be required. The high spatial resolution of confocal microscopy generates three-dimensional renderings of entire body parts, allowing us to define the position of structures relative to tissues that have straightforward homology assignments (Klaus, Kulasekera, & Schawaroch, 2003). Many developing tissues progressively become more complex over developmental time. The formation of specific traits is often established only after the tissue that encompasses that trait is identifiable, providing clear anchor points in a conserved tissue to establish homology. Thus, developmental trajectories provide a rich context in which to disentangle ambiguous relationships among rapidly evolving structures.

The terminalia (genitalia and analia) of drosophilids host an extensive assortment of rapidly evolving morphological structures. Variation of terminal structures is often one of the most reliable ways to differentiate species of *Drosophila* based on adult morphology (Bock & Wheeler, 1972; Hsu, 1949; Markow & O’Grady, 2006; Okada, 1954). The male genital structures are often divided into two major compartments: the periphallic parts surrounding the anus, which mostly play a role in grasping the external surface of the female genitalia (Acebes, Cobb, & Ferveur, 2003; Jagadeeshan & Singh, 2006; Kamimura & Mitsumoto, 2011; Masly & Kamimura, 2014; Mattei, Riccio, Avilaa, Wolfner, & Denlinger, 2015; Rhebergen, Courtier-Orgogozo, Dumont, Schilthuizen, & Lang, 2016; Robertson, 1988; Yassin & Orgogozo, 2013), and the phallic parts (Figure 1), which comprise the intromittent organ. While the homology of the periphallic organs has always been relatively straightforward, the phallus itself has posed several challenges in this regard. In particular, the homology of the various phallic processes, pointed outgrowths, remains controversial (Figure 2, Supplementary videos) (reviewed in Rice et al., 2019). These outgrowths have been implicated in sexual conflict between males and females, and in some species have been shown to physically interact with corresponding pockets in the female genitalia (Kamimura, 2016; Muto, Kamimura, Tanaka, & Takahashi, 2018; Yassin & Orgogozo, 2013) or induce copulatory wounds (Kamimura, 2007). Male seminal proteins are associated with increased ovulation and reduced remating rates and can enter the female circulatory system through these copulatory wounds (Avila, Sirot, Laflamme, Rubinstein, & Wolfner, 2011; Kamimura, 2010). To better understand how genital morphology may coevolve we must better establish which homologous tissues have been modified in each sex.

**Figure 1:**
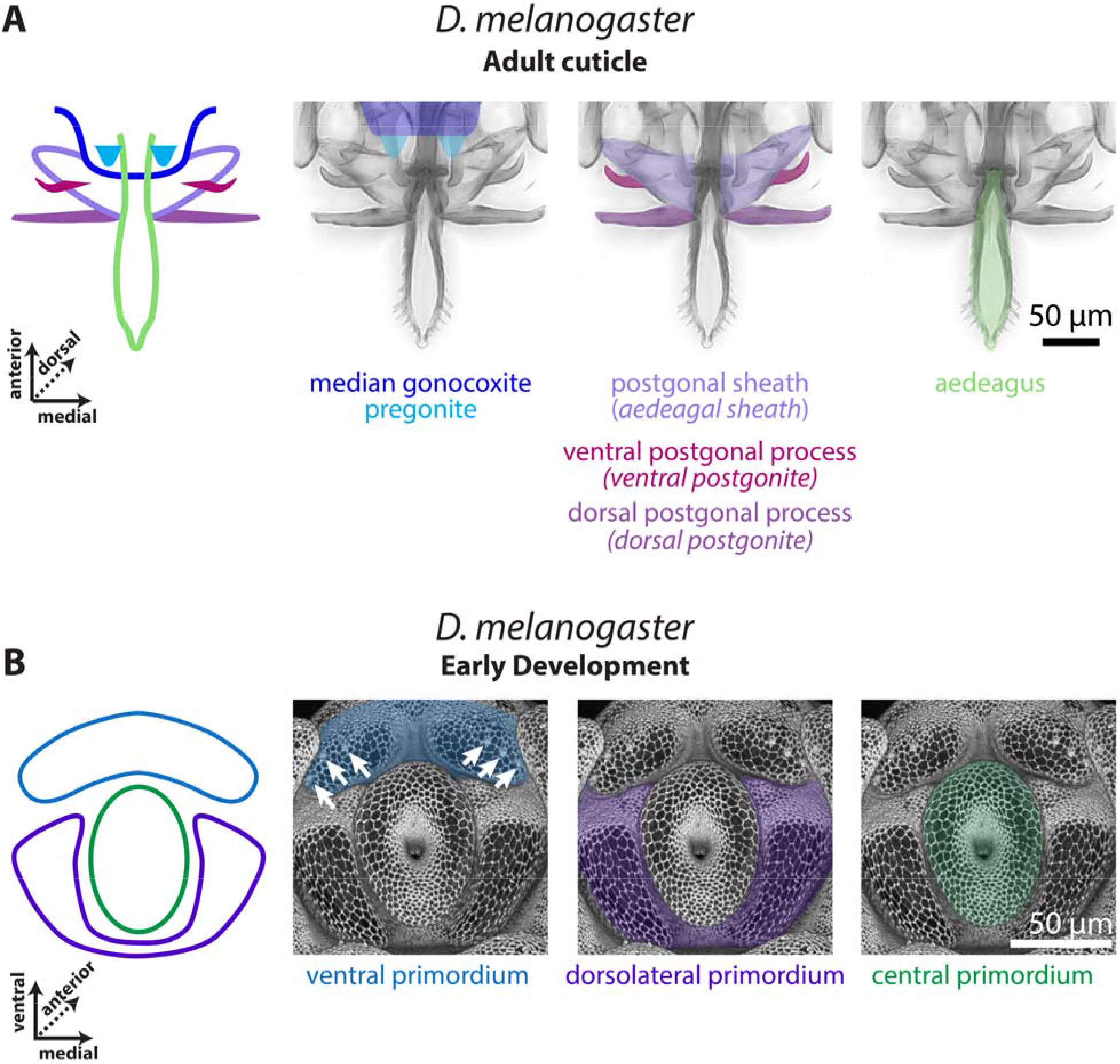
The *D. melanogaster* pupal phallus is produced by three primordia **A)** Left: A schematic representation of the adult phallus of *D. melanogaster*, the median gonocoxite outlined in dark blue, the pregonites highlighted in light blue, the postgonal sheath in light purple, the ventral postgonal process in magenta, the dorsal postgonal process in violet, and the aedeagus in light green. Right: the adult phallus of a *yw;+:+* line of *D. melanogaster*. **B)** Left: A schematic representation of the early developing pupal genitalia of *D. melanogaster*. The primordia of developing phallus with the ventral primordium in blue, the dorsolateral primordium in purple, and the central primordium in green. Right: Developing pupal phallus of a *D. melanogaster* arm-GFP transgenic line. Apical cellular junctions are shown, highlighting the overall morphology. White arrows indicate the position of the pregonal bristles. Aedeagal sheath is an alternative term for postgonal sheath, ventral postgonite is an alternative term for ventral postgonal process, and dorsal postgonite is an alternative term for dorsal postgonal process. The images of the developing phallus are shown with ventral view on top to match the orientation of the phallus during copulation.

**Figure 2:**
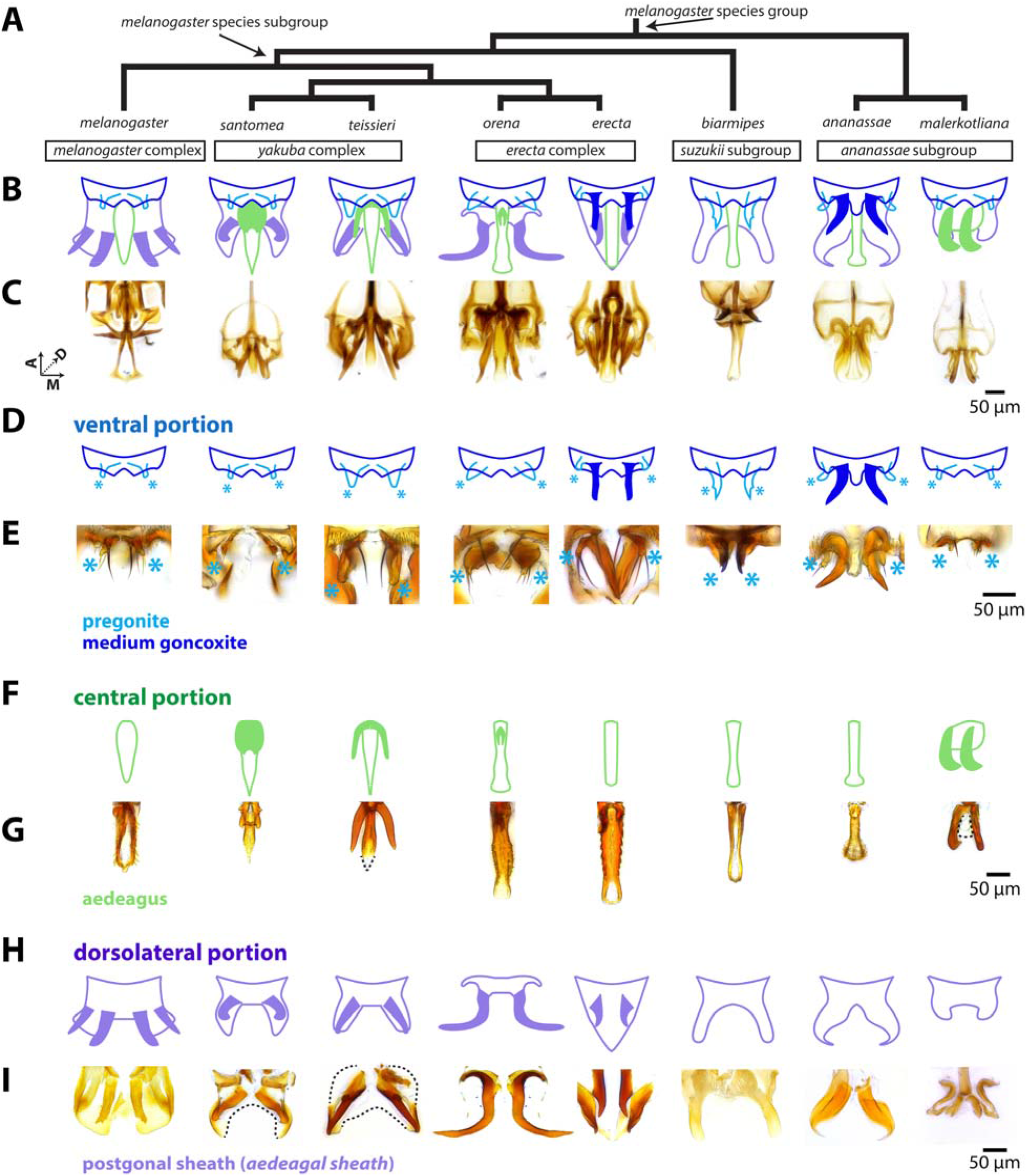
The rapidly evolving phallus is composed of three main components **A)** Phylogeny for eight species of the *melanogaster* species group based on Obbard et al., 2012 with nodes that contain the *melanogaster* species subgroup and *melanogaster* species group indicated by arrows. **B)** A schematic breakdown of the adult phalluses of each species. **C)** Light microscopy images of the whole adult phallus for each species. Image stacks that show the relative position of each part can be found in Supplementary videos. **D)** Schematic representation of the ventral portion of the phallus (dark blue) which contains the pregonites (light blue) and processes (filled in dark blue) in *D. erecta* and *D. ananassae*. Light blue asterisks designate the position of the pregonites. **E, G, I**: Light microscopy images of microdissections of the adult phallus (here in lateral view, distal end pointing downward) separated in the ventral, central, and dorsolateral portions. **E)** Microdissections of the ventral portion processes shows the processes of *D. erecta* and *D. ananassae* are connected to the pregonites. Light blue asterisks designate the position of the pregonites. **F)** Schematic representation of the central portion of the phallus (light green), which contains processes (filled in light green) in *D. santomea, D. teissieri, D. orena*, and *D. malerkotliana*. **G)** Microdissections of the aedeagus confirm that the processes are physically attached to aedeagus. The aedeagus of the *D. malerkotliana* is translucent (dashed line) with only the process sclerotized. **H)** Schematic representation of the dorsolateral portion of the phallus (light purple), which contains two pairs of processes (filled in light purple) in *D. melanogaster*, and one pair in *D. santomea, D. teissieri, D. erecta, and D. orena*. **I)** Microdissection confirms that the processes are physically attached to postgonal sheath. In *D. santomea, D. teissieri, D. erecta*, and *D. malerkotliana* portions of the anterior postgonal sheath are translucent (outlined with dashed lines). The image of the *D. orena* dorsolateral portion was created by copying and mirroring one side of the structure, as it was difficult to flatten intact for imaging. Aedeagal sheath is an alternative term for postgonal sheath.

Two major sources have contributed to confusion regarding homology of phallic processes. The first is the relationship of the postgonal sheath (referred to as aedeagal sheath in Rice et al., 2019) with respect to these processes (Figure 2 H-I). Several authors consider the postgonal sheath and postgonal processes (referred to as postgonites in Rice et al., 2019) of *D. melanogaster* as substructures of a unified tissue that was usually referred to as the “posterior parameres” (Bock & Wheeler, 1972; Okada, 1954; Tsacas, Bocquet, Daguzan, & Mercier, 1971). While others designated the postgonal sheath as a separate tissue from the postgonal processes (Al Sayad & Yassin, 2019; Bächli, Vilela, Andersson Escher, & Saura, 2004; Lachaise et al., 2004). The three-dimensional nature and the presence of transparent cuticle has made it difficult to determine the precise connection points of the processes to the tissues of the phallus. Determining whether these processes were formed by a single or separate primordium would help resolve this discordance. The second source of confusion is in regard to the nomenclature used to compare the phallic processes in different members of the *melanogaster* species group. The term “basal process” has been used to refer to a number of pointed outgrowths that are attached to different phallic tissues in different species (Kamimura, 2007, 2010, 2016; Kamimura & Mitsumoto, 2011, 2012a, 2012b; Kamimura & Polak, 2011). Such a designation implies a concept of homology independent of the exact anatomical position. Yassin & Orgogozo (2013) sought to provide distinct terms, such as “spurs” and “hooks” for outgrowths emanating from the same tissue, implying the potential for non-homology. Building upon our recent revision of the male terminalia nomenclature of *D. melanogaster* (Rice et al., 2019), developmental studies presented here allow us to provide a more detailed, homology-informed nomenclature for these structures.

In this study, we characterized both the adult morphology and the development of the pupal genitalia in five members of the *melanogaster* species subgroup and three outgroup members from the larger *melanogaster* species group. This analysis allows us to determine whether processes are homologous or of different origins. Tracing the development of the phallus by confocal microscopy showed that all processes arise from three distinct pupal regions that likely represent primordia for three subdivisions of the phallus in all species. We found both that several similarly shaped processes arise from distinct primordia, whereas in other cases, distinct processes arise from different parts of the same primordium. In light of these analyses, we refined the identity and terminology of the phallic processes and identify distinct homology groups. We map these different morphologies on previously established phylogenies and identified multiple gain, loss, and homoplastic events in the history of these diverse structures. Thus, our results demonstrate how developmental approaches can resolve unclear relationships among rapidly evolving structures.

## Materials and Methods

### Drosophila strains

To study the evolution of the phallic processes in the *melanogaster* species subgroup we used the following species, representing all major complexes: *D. santomea* (Lachaise et al., 2000), *D. teissieri* (Tsacas, 1971), *D. orena* (Tsacas & David, 1978), *D. erecta* (Tsacas & Lachaise, 1974), *D. melanogaster* (Meigen, 1830) from the *melanogaster* species subgroup and the following outgroup species *D. biarmipes* (Malloch, 1924) from the *suzukii* subgroup, *D. ananassae* (Doleschall, 1858), and *D. malerkotliana* (Parshad & Paika, 1964) from the *ananassae* subgroup. Previous work has investigated the function of the copulatory anatomy of all species we analyzed (Kamimura, 2007, 2016; Muto et al., 2018; Yassin & Orgogozo, 2013). Stocks were obtained from both the National Drosophila Species Stock Center at Cornell (*D. santomea* (14021-0271.01), *D. teissieri* (14021-0257.01), *D. orena* (14021-0245.01), *D. erecta* (14021-0224.01), *D. biarmipes* (14023-0361.09), *D. ananassae* (14024-0371.13), the Bloomington Drosophila Stock Center *D. melanogaster* armadillo-GFP, arm-GFP, (Bloomington stock number #8556), and from the lab of Dr. Thomas Williams, *D. malerkotliana*.

### Sample collection, dissection, and fixation

Male white pre-pupae were collected at room temperature and incubated in a petri dish containing a moistened Kimwipe at 25°C prior to dissection. After incubation, pupae were impaled in their anterior region and immobilized within a glass dissecting well containing Phosphate Buffered Saline (PBS). The posterior tip of the pupa (20-40% of pupal length) was separated and washed with a P200 pipette to flush the pupal terminalia into solution. Samples were then collected in PBS with 0.1% Triton-X-100 (PBT) and 4% paraformaldehyde (PFA, E.M.S. Scientific) on ice, and multiple samples were collected in the same tube. Samples were then fixed in PBT + 4% PFA at room temperature for 30 min, washed three times in PBT at room temperature, and stored at 4°C.

### Immunohistochemistry and microscopy

After fixation developing pupal genitalia of all species except *D. melanogaster* were stained with rat anti-E-cadherin, 1:100 in PBT (DSHB Cat# DCAD2,RRID:AB_528120) overnight at 4°C, followed by an overnight at 4°C incubation with anti-rat 488,1:200 (Invitrogen, Carlsbad, CA) to visualize apical cell junctions. For *D. melanogaster*, an *armadillo-GFP* tagged line (Bloomington stock number #8556) was used to visualize apical cell junctions (Huang et al., 2012). Fluorescently labeled samples were mounted in glycerol mounting solution (80% glycerol, .1M Tris, pH 8.0) on microscope slides coated with poly-L-lysine (Thermo Fisher Scientific #86010). Samples for all species except *D. melanogaster* were imaged at 20X on a Leica TCS SP8 confocal microscope. *D. melanogaster* samples were imaged at either 20X or 40X on an Olympus Fluoview 1000. As the imaged structures are three-dimensional in nature, we used the MorphoGraphX program (de Reuille et al., 2015) to render and manipulate images in three-dimensions. This allowed us to rotate the samples to better present the most informative perspectives of the various phallic structures.

For light microscopy of adult phallic microdissections, samples were mounted in PVA Mounting Medium (BioQuip) until fully cleared and imaged at 20X magnification on a Leica DM 2000 with a Leica DFC450C camera and the resulting images were enhanced using Adobe Photoshop. For light microscopy images and videos of the whole phallus, genitalia were dissected in water, cleared overnight in 10% KOH at RT. These were them mounted in a drop of Dimethyl Hydantoin Formaldehyde (Steedman, 1958) on a coverslip and oriented using 2 mounting needles before the resin hardened. Coverslip were positioned on a microscope slide, the hard drop facing away from the microscope lens. Images were acquired on Ti2-Eclipse Nikon microscope equipped with a 20x plan apochromatic lens and a 5.5 M sCMOS camera (PCO, Kelheim, Germany). Each preparation was imaged as a z-stack (z-step = 2 μm). The stacks are presented as raw images. Stacks of images were also projected into single extended depth-of-field images using Helicon Focus software (HeliconSoft) and the resulting projections were enhanced using Adobe Photoshop.

### Establishing landmarks for early, middle, and late timepoints

We used confocal microscopy to chart a time course of the developing phallus (Figures S1-S3). To compare the development of the phallus of these species, we needed to examine whether all analyzed species develop at the same rate after pupal formation. Due to the large-scale changes in the phallus of these species, we used two stable features found outside of the phallus to calibrate developmental timing. In all analyzed species, the epandrial ventral lobe (lateral plate) and surstylus (clasper) first appear as a single continuous structure early in development, but then separate from each other as development progresses (Figure S2). We use the beginning of this separation as a landmark for the “early” developmental timepoint. We also used the midpoint of this progression to approximate the “mid” timepoint. This intermediate timepoint is useful in showing which tissue the phallic processes protrude from during development. In all species, the adult cerci (anal plates) directly abut against one another but during “early” and “mid” development, these structures are separated from one another by a large gap (Figure S3). We designate “late” timepoint as directly preceding the closing of this gap between the cerci.

## Results

### Unpigmented cuticle reveals undescribed connection points in the phallus

In order to better understand how the processes surrounding the aedeagus are physically connected to the neighboring tissues of the phallus, we imaged whole (Figure 2 B,C) and micro-dissected adult phalluses in eight members of the *melanogaster* species group (Figure 2 D-I). The phallus of each species can be partitioned into three discrete parts. The ventral portion (Figure 2 D,E) contains the pregonites, an outgrowth that contains three bristles, and the median gonocoxite (the central section of the shield shaped hypandrium). The central portion (Figure 2 F,G) contains the aedeagus, through which sperm is transferred. The dorsolateral portion contains the postgonal sheath (referred to as aedeagal sheath in (Rice et al., 2019)), a flat sheet that wraps around the aedeagus, and the pair of processes known as the postgonal processes (referred to as postgonites in Rice et al., 2019) (Figure 2 H,I). Analysis of these dissections support the designation of the postgonal sheath and postgonal processes as a single tissue (Bock & Wheeler, 1972; Okada, 1954; Tsacas et al., 1971). Furthermore, we found that certain species had processes connected to different portions of the phallus— a ventral portion (*D. erecta, D. ananassae*), a central portion (*D. santomea, D. teissieri, D. orena*, and *D. malerkotliana*), and a dorsolateral portion (all members of the *melanogaster* subgroup).

While imaging, we observed that parts of the postgonal sheath in the *melanogaster* species subgroup and *D. malerkotliana* were partially translucent, and only detectible after microdissection. It is this translucent tissue of the postgonal sheath that physically connects to the postgonal processes in *D. melanogaster* (Figure2l). These observations highlight that, due to their transparency, determining the exact connection points between the processes and the rest of the phallus can be difficult to visualize by traditional light microscopy approaches. To test whether the different connection points of the phallic processes observed in the adult reflect separate homology groups we investigated whether these phallic processes were initially produced by the same or different primordia during development.

### Phallic structures develop from three primordia in D. melanogaster

To date, the morphogenesis of the three-dimensional adult phallic structures from the epithelium of the larval genital disc has been investigated only in *D. melanogaster* (Ahmad & Baker, 2002; Epper, 1983). Additionally, using surgical fragmentation of the larval genital disc, Bryant & Hsei, 1977 provided a fate map for the different adult structures. They showed that the phallus is situated at the subcenter of the symmetrical imaginal disc and is surrounded on each side by a primordium that will produce the medium gonocoxite and pregonites. However, the sequence and timing of the appearance of the various substructures of the phallus during development, remains unknown. By finding the key points in development where substructures first emerge, we can determine the primordium from which each process initially forms.

Early in *D. melanogaster* pupal development (see timepoint determination in the Materials and Methods) the phallus is separated into three stereotypic regions that likely represent primordia: ventral, dorsolateral and central (Figure 1B). As the pupal phallus continues to develop from this point, the ventral primordia form a pair of processes (Figure 1A). This pair develops into the small processes known as the pregonites that can be recognized from the presence of minute bristles (Figure 1B arrows), while the remainder of the primordia forms the median gonocoxite (Figure S4). The dorsolateral primordia produce two processes (one dorsal and one ventral) (Figure 3B). These processes ultimately develop into the ventral and dorsal postgonal processes (referred to as ventral postgonite and dorsal postgonite in Rice et al., 2019) (Figure 1A) The remaining parts of the dorsolateral tissue develop into the large flaps of the postgonal sheath (Figure 1A, Figure S5). The central primordium of *D. melanogaster* develops into the aedeagus and lacks a process (Figure 1A, Figure S6).

**Figure 3:**
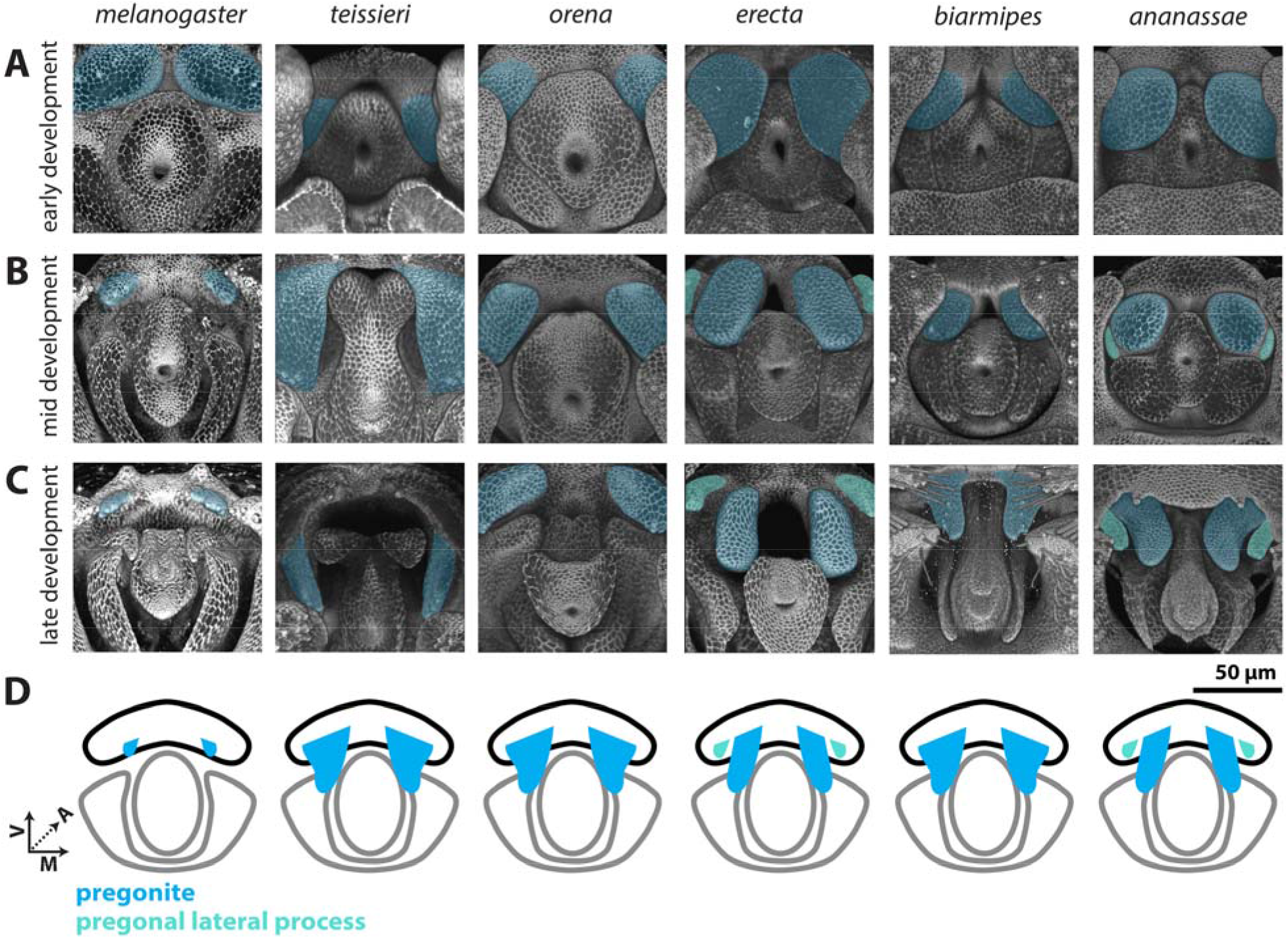
Processes developing from the ventral primordium are found in all members of the *melanogaster* species group. **A-C)** Signal from the apical cellular junctions was used to highlight the overall morphology of developing pupal genitalia. *D. teissieri, D. orena, D. erecta, D. biarmipes*, and *D. ananassae* were stained for ECAD while apical cell junctions were visualized in *D. melanogaster* by detecting *arm-GFP* (see methods). **A)** Early in development, a pair of processes, that will form the pregonites, can be visualized in the ventral primordia in all species shown (light blue). **B)** By mid-development, large processes can be found in all species shown except *D. melanogaster*. In *D. erecta* and *D. ananassae*, the pregonite is split into a large pregonal medial process and a small lateral bristle-bearing process (teal). **C)** By late development, the pregonites have extended to their full adult size and shape. The pregonites are connected to the medial-ventral portion of the median gonocoxite, see Figure S4. **D)** Schematic representations of the pregonites (blue) and pregonal lateral process (teal) showing their approximate size, number, and connections to the medial gonocoxite (outlined in black). All images have the same axes, V (Ventral), A (Anterior), M (Medial) and are the same scale.

### The three primordia are conserved across species

Several studies have analyzed pupal development of the terminalia in species outside of *D. melanogaster*, but did not investigate phallic structures (Glassford et al., 2015; Hagen et al., 2019; Smith, Davidson, & Rebeiz, 2020). To determine whether the features of phallic development observed in *D. melanogaster* are conserved in members of the *melanogaster* species group, we produced a developmental time course for the remaining seven species studied here (Figure S1-S3). Our time course indicates that all adult phallic organs develop from three regions that are similar in size and shape to the ones described in *D. melanogaster* and thus likely represent homologous primordia. The ventral, dorsolateral, and central primordia produce the median gonocoxite (Figure S4), pregonal sheath (Figure S5), and aedeagus (Figure S6), respectively in all species. Nonetheless, significant interspecific differences were observed regarding the timing of development (Figure S1). As we only used one strain per species, we cannot comment if these are particular properties of the strains/laboratory conditions we used or are general differences between the species. We found that most species had early, mid, and late developmental timepoints within a six-hour window of each other (Figure S1-S3).

### Different processes emerge from different primordia

The developmental analysis of the eight species used in this study allowed us to test whether the phallic processes seen in the adults of each species were produced by the same primordia. We began by investigating the ventral primordium (Figure 3, Figure S4), which develops into the pregonites in all analyzed species. While the size of the pregonites varies between species, during mid-development (Figure 4B) we can identify recognizable outgrowths from the ventral primordium, consistent with a highly conserved developmental trajectory. Interestingly, an additional pregonal process is found in two distantly-related species, *D. erecta* and *D. ananassae*. Both *D. erecta* and *D. ananassae*, produce two processes from their ventral primordia, a large pregonal medial process and a second smaller pregonal lateral process which contains the three pregonal bristle cells (Figure S7) and overall resembles the pregonites of other species. To determine whether this additional process was produced by duplication or fission of the pregonite we inspected early pupal timepoints. We found that initially a single process is formed (Figure 3A), which during mid-development asymmetrically splits along the medial-lateral axis to form the distinct lobe-like pregonal medial process (Figure 3B). These asymmetric projections then extend in late development to form the larger pregonal medial process and smaller pregonite (Figure 3C). Thus, although the ventral primordium produces the pregonite in all species we examined, in *D. erecta* and *D. ananassae* the ventral primordium is split into the pregonite and a pregonal medial process.

**Figure 4:**
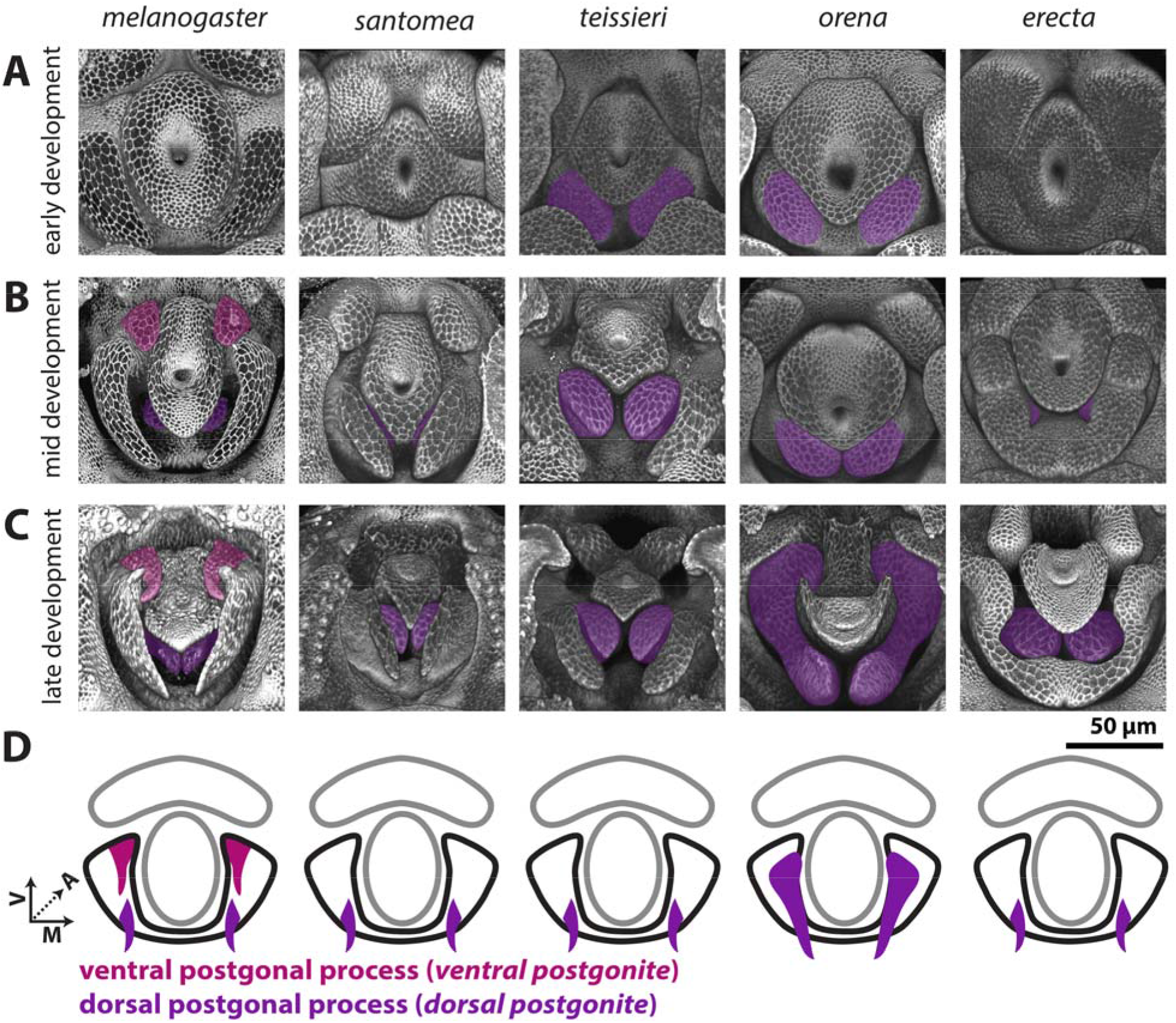
Processes produced by the dorsolateral primordium are found only in the *melanogaster* subgroup. **A-C)** Signal from the apical cellular junctions was used to highlight the overall morphology of developing pupal genitalia. *D. santomea, D. teissieri, D. orena*, and *D. erecta*, were stained for ECAD while *D. melanogaster* samples used arm-GFP. **A)** Early in development, the dorsolateral primordium is a smooth lobe like structure in all analyzed species. **B)** By mid-development, all shown species form processes in the dorsal portion of the dorsolateral primordium (violet). *D. melanogaster* also forms an additional pair of processes in the ventral portion of dorsolateral primordium (magenta). **C)** By late development, the dorsal and ventral processes have extended to a long thin shape. Both the ventral and dorsal processes are connected to the medial-anterior part of the postgonal sheath which is formed by the remaining tissue of the dorsolateral primordium. **D)** Schematic representations of the dorsal (violet) and ventral (magenta) postgonal process showing where they connect to the postgonal sheath (outlined in black). All images have the same axes, V (Ventral), A (Anterior), M (Medial) and are the same scale. Ventral postgonite is an alternative term for ventral postgonal process, and dorsal postgonite is an alternative term for dorsal postgonite (Table 1).

The dorsolateral primordium (Figure 4, Figure S5) showed a number of large evolutionary changes within the *melanogaster* species group. We found that no species, other than *D. melanogaster*, develop the ventral process that forms the ventral postgonal process (Figure 4). By contrast, all members of the *melanogaster* species subgroup form dorsal postgonal processes. Outside of the *melanogaster* subgroup, we did not find any modifications of the dorsolateral primordium, which develops into a single thin, strongly sclerotized structure in those species that resembles the postgonal sheath of *D. melanogaster*. However, the size, and shape of these homologous structures significantly differ among species (Figure 2 H,l), ranging from the flat rod-like processes in *D. biarmipes*, to the strongly pointed sinuate processes in *D. ananassae*, and the minute, transparent sclerites in *D. malerkotliana*.

While the central primordium (Figure 5, Figure S6) forms a simple aedeagus that lacks processes in *D. melanogaster, we* note processes which develop in *D. santomea, D. teissieri, D. orena* and *D. malerkotliana*. Early in development, the central primordia of all species analyzed are similar in size and shape (Figure 5A). However, during mid-development, in *D. santomea, D. teissieri*, and *D. orena*, the ventral side of the central primordium elongates to form a process (Figure 5B). The process of *D. teissieri* and *D. orena* splits along the ventral midline to form a pair of processes, while in *D. santomea*, it forms one rounded structure. These processes further elongate in late development to more closely resemble the size and shape of their adult counterparts (Figure 5C).

**Figure 5:**
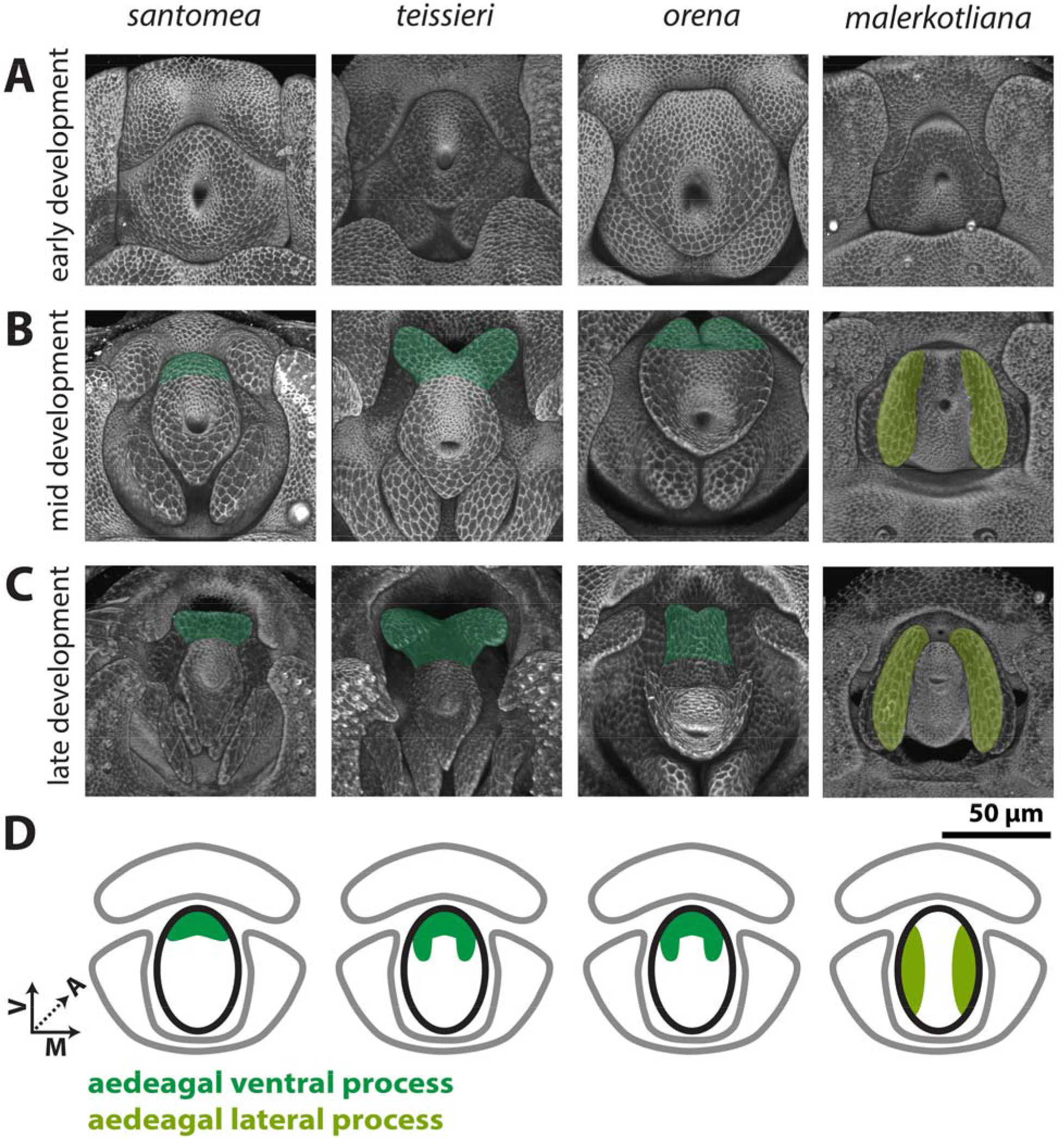
Processes developing from the central primordium are found in the *yakuba/erecta* and *bipectinata* complexes. **A-C)** Signal from the apical cellular junctions (ECAD) was used to highlight the overall morphology of developing pupal genitalia. **A)** Early in development, the central primordium forms a flat donut-shaped structure in all species shown. **B)** By mid-development, the ventral portion of the aedeagus is extended in *D. santomea, D. teissieri*, and *D. orena* in what will form the aedeagal ventral process (dark green). In *D. malerkotliana* the lateral edges of the central primordium extend anteriorly in what will form the aedeagal lateral process (yellow-green). **C)** By late development, the aedeagal ventral process and the aedeagal lateral process further extend from the aedeagus. **D)** Schematic representations of the aedeagal ventral process (dark green) and aedeagal lateral process (yellow-green) showing where they connect to the aedeagus (outlined in black). All images have the same axes, V (Ventral), A (Anterior), M (Medial) and are the same scale.

Okada, 1954 and Bock & Wheeler, 1972 suggested that the aedeagus in the *melanogaster* species group were of two types: fused like in *D. ananassae* and split like in *D. malerkotliana*. Indeed, we did not observe any process in the central primordium of *D. ananassae*, whereas a pair of processes develops in *D. malerkotliana*. Kamimura, 2007 suggested that the aedeagus of *D. malerkotliana* has degenerated and was replaced by a pair of lateral processes. During early development, the central primordium of *D. malerkotliana* is similar to all other analyzed species (Figure 5A). However, by mid-development, the lateral sides of the central primordium extend, forming a pair of processes, while the medial-dorsal and medial-ventral sides of the central primordium fail to extend (Figure 5B). Late in development, the proximal-dorsal side of the lateral process constricts, conferring a hook like shape (Figure 5C). As this substructure is produced from the lateral portions of the central primordium and not from the ventral portion, it is likely non-homologous to the aedeagal ventral processes of the *yakuba* and *erecta* complexes. We therefore propose the term aedeagal lateral process for this substructure of *D. malerkotliana*.

## Discussion

The rapid evolution of morphological structures is an attractive subject for study, as it allows us to glimpse at the molecular and genetic causes of remodeled and restructured anatomical forms. Here, we examined some of the most rapidly evolving morphologies of *Drosophila melanogaster* and its close relatives. Despite decades of research, many of the intricate phallic processes have eluded our ability to clearly classify their homology relationships. By studying the developmental trajectories of these processes in multiple species, we have better defined their physical connections, and clarified which structures most likely share ancestry. This research highlights the distinct challenges in studying novelties at mesoevolutionary scales, specifically that traits may be rapidly gained and lost between closely related species making it difficult to discern between true homology and convergence (Abouheif, 2008).

### Classification and nomenclature of rapidly evolving phallic structures

Our results suggest that the great diversity of the phallic structures of the eight species studied here cluster into three homology groups corresponding to the three pupal primordia, leading us to propose revised naming conventions. First, our developmental analysis supports the notion, initially suggested by Okada, 1954, that the weakly sclerotized postgonal sheath and strongly sclerotized postgonal processes in *D. melanogaster*, are both parts of the same tissue, which *Drosophila* systematists called the “posterior paramere” e.g. (Bock & Wheeler, 1972; Tsacas et al., 1971). Because the term “posterior paramere” is itself synonymous to the term “postgonite” in Dipteran systematics (Tsacas et al., 1971; van Emden & Hennig, 1970), we suggest using the term “postgonite” to encompass the combined tissue produced by the dorsolateral primordia in species of the *melanogaster* group (including both the postgonal sheath and the processes), and the term “postgonal processes” to designate the strongly sclerotized branches emerging from this tissue in the *melanogaster* subgroup. Second, our results also show that the structures previously called the “basal processes” (Kamimura, 2007, 2010, 2016; Kamimura & Mitsumoto, 2011, 2012a, 2012b; Kamimura & Polak, 2011), develop from different primordia and are therefore most likely non-homologous. We suggest therefore to give them distinct names that directly relate to the tissues that produce them: aedeagal ventral process in species of the *yakuba* complex (synonymous to Yassin & Orgogozo, 2013 phallic spur) and *D. orena* (synonymous to Yassin & Orgogozo, 2013 phallic hook), the pregonal medial process in *D. erecta* and *D. ananassae*, and aedeagal lateral process in *D. malerkotliana* and species of the *bipectinata* complex (Table 1). Future work that establishes the extent of cell migration between or with the three regions that we designate as primordium will further improve our resolution of the homology of these phallic processes.

**Table 1:** Table of correspondence between terms previously used in publications and proposed nomenclature.

|  |  |  |
| --- | --- | --- |
| 843 | **Tables:** |  |
|  | **New nomenclature** | **Previous terminology** |
|  | postgonite | posterior paramere (Bock & Wheeler, 1972; Tsacas et al., 1971) |
|  | dorsal postgonal process | dorsal postgonite (Rice et al., 2019; Vincent et al., 2019)<br>dorsal branch (Kamimura, 2010, 2016; Kamimura & Mitsumoto, 2011)<br>dorsal paramere (Bryant & Hsei, 1977) |
|  | ventral postgonal process | ventral postgonite (Rice et al., 2019; Vincent et al., 2019)<br>ventral branch (Kamimura, 2010; Kamimura & Mitsumoto, 2011)<br>ventral paramere (Bryant & Hsei, 1977) |
|  | aedeagal ventral process | phallic spur (Yassin & Orgogozo, 2013)<br>phallic hook (Yassin & Orgogozo, 2013)<br>ventral branch (Kamimura, 2012, 2016; Kamimura & Mitsumoto, 2012b, 2012a; A. E. Peluffo et al., 2015; A. Peluffo et al., 2021) |
|  | aedeagal lateral process | basal process (Kamimura, 2007; Kamimura & Polak, 2011) |
|  | pregonal medial process | basal process (Kamimura, 2007), ventral branch (Kamimura, 2016) |
|  | Postgonal sheath | Aedeagal sheath (Rice et al., 2019; Vincent et al., 2019) |
| 844 | Table 1: Table of correspondence between terms previously used in publications and proposed |  |
| 845 | nomenclature. |  |

### Evolution of the phallic structures

Mapping character states over robust phylogenies provide the opportunity to distinguish novel from recurrent (homoplastic) states as well as derived states (synapomorphic) from ancestral (symplesiomorphic) ones. Our findings have led us to propose a model for the evolution of the phallic processes found in the *melanogaster* species subgroup (Figure 6). For example, our demonstration of the development of an additional pregonal process in *D. erecta* and *D. ananassae* (Figure 3, S4) is likely recurrent, as illustrations from Bock & Wheeler, 1972 suggest that this configuration of the pregonites might have recurrently evolved in this clade. Reversals to ancestral states through secondary losses represent another mechanism of recurrent evolution. The lack of the aedeagal ventral processes in *D. erecta*, is more likely due to loss rather than an independent gain of the aedeagal ventral process in *D. orena*.

**Figure 6:**
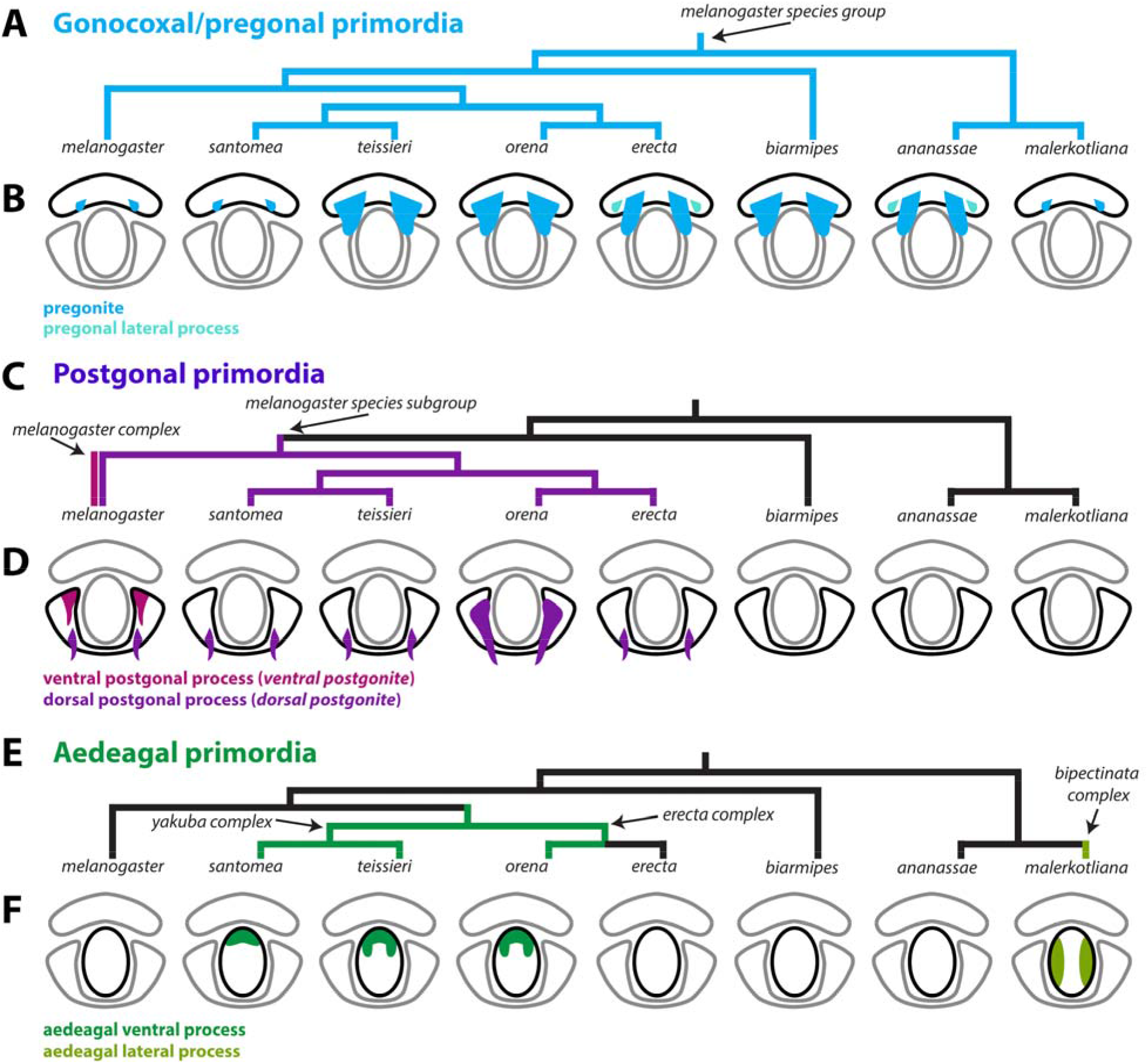
A model of the evolution of phallic processes in the *melanogaster* species group **A,C,E)** Phylogeny of the 8 analyzed species based on Obbard et al., 2012. **A)** Parsimony suggests that the pregonites originated outside of the *D. melanogaster* species group. **B)** Light blue represents the pregonites and teal represents the pregonal lateral process. **C)** Parsimony suggests that the ventral postgonites originated in the *melanogaster* complex (*D. melanogaster, D. simulans, D. mauritiana, D. sechellia*) and that the dorsal postgonal process originated in the *melanogaster* subgroup. D) Magenta represents the ventral postgonal process, violet represents the dorsal postgonal process. **E)** Parsimony suggests that the aedeagal ventral process originated at the base of the *erecta* and *yakuba* complexes. Additionally, parsimony suggests that the aedeagal lateral process originated in the bipectinata complex. **F)** Dark green represents the aedeagal ventral process and yellow-green represents the aedeagal lateral process. Ventral postgonite is an alternative term for ventral postgonal process, and dorsal postgonite is an alternative term for dorsal postgonite. Future developmental analysis across more members within and outside of the *D. melanogaster* species group will be required to better establish when these distinct phallic processes first originated.

In the *melanogaster* subgroup, all species contain a strongly-sclerotized dorsal postgonal process which develops as a localized extension within the dorsolateral primordium. The development of a strongly-sclerotized ventral postgonal process is a definitive novelty in *D. melanogaster* and allied species of the *melanogaster* complex. Although we did not find structures resembling the dorsal postgonal processes in members outside of the *melanogaster* species subgroup that we studied here, polarization remains difficult. Indeed, Okada, 1954 and Bock & Wheeler, 1972 reported the presence of “basal processes of the posterior parameres” in multiple members of the *melanogaster* species group. Similarly, Bächli et al., 2004 illustrated the presence of “ribbon-shaped process” in several members of the *obscura* group which is sister to the *melanogaster* species group. Further taxonomic sampling and better phylogenetic resolution of those clades are required to draw a more complete picture of the evolution of the postgonal differentiation outside the *melanogaster* subgroup. The novel structures described here may present an excellent model to study the molecular mechanisms producing novelty.

Although we do not address the function of the phallic processes, other research groups have demonstrated that the rapid evolution of male genital sclerites in arthropods were most likely driven by selection (Eberhard, 1985; Hosken, Archer, House, & Wedell, 2019; Simmons, 2014). These included groups as diverse as spiders (Huber, 2005), damselflies (Cordero-Rivera, 2017), waterstriders (Rowe & Arnqvist, 2012), moths (McNamara, Dougherty, Wedell, & Simmons, 2019) and beetles (Simmons & Fitzpatrick, 2019). This unique mechanism is remarkable given the diverse developmental origin of the rapidly evolving male structures in these groups, being pedipalps in spiders (Quade et al., 2019), cerci in damselflies (McPeek, Shen, & Farid, 2009), pregenital segments in waterstriders (Perry & Rowe, 2018), chitinous spermatophores in butterflies (Sánchez & Cordero, 2014), or aedeagii of appendicular and non-appendicular origins in bugs and beetles (Aspiras, Smith, & Angelini, 2011). However, the exact form of selection, e.g., cryptic female choice, sexual conflict, etc., remains debated in the literature (Ah-King, Barron, & Herberstein, 2014; Brennan & Prum, 2015; Eberhard, 1985; Masly, 2012).

In Drosophila, the various processes of the phallus have been implicated in copulatory wounding of the female (Kamimura, 2007, 2016; Muto et al., 2018; Yassin & Orgogozo, 2013). Furthermore, studies have found that some of the phallic processes pivot from pointing posteriorly to pointing laterally, when the phallus is everted during copulation, thus directing how they interact with the female reproductive tract (Kamimura, 2010). The ability to pivot during copulation correlates with the homology groups we have found in this study. The ventral and dorsal postgonal processes, and pregonites pivot during copulation while the aedeagal ventral process does not change orientation. This may be due to the direct connection of the aedeagal ventral process to the aedeagus. Surprisingly the aedeagal lateral process, which is also directly connected to the aedeagus, pivots laterally during copulation, which may only be possible due to the loss of aedeagal sclerotization, making the tissue between the aedeagal lateral processes flexible. Co-evolution between the phallic processes and the female genitalia has been suggested and several novel modifications of the female genitalia have been identified (Kamimura, 2007; Yassin & Orgogozo, 2013). A developmental analysis of the female genitalia of these species along with three-dimensional analysis of copulating flies similar to studies in *D. melanogaster* (Mattei et al., 2015; Shao et al., 2019) would provide vital context for the potential co-evolution of novel male and female structures.

### Developmental mechanisms underlying phallic evolution

A major challenge in the evo-devo field has been to identify the molecular mechanisms driving morphological novelty (Linz, Hu, & Moczek, 2020; Moczek, 2008; Rebeiz, Patel, & Hinman, 2015; G. P. Wagner & Lynch, 2010). While macroevolutionary novelties have been the focus of coarse-grained molecular study (Bruce & Patel, 2020; Clark-Hachtel & Tomoyasu, 2020; Emlen et al., 2006; Hinman et al., 2003; Prud’Homme et al., 2011), much hope has been placed on rapidly diverging structures in molecularly amenable systems (Rebeiz & Tsiantis, 2017). Recent work in Drosophila genital evolution has highlighted how quickly changes in cellular morphology (Green et al., 2019; Smith et al., 2020), and genetic networks (Glassford et al., 2015; Hagen et al., 2021; Nagy et al., 2018) can lead to shifts in morphology in closely related species. Our work highlights distinct underexplored challenges to interpreting and advancing these model systems. The ambiguous ancestry of similar parts which appear in different locations causes us to consider multiple models to explain their emergence. These parts may arise by parallelism - a predisposition to drive similar new structures by co-opting the same networks (Abouheif, 2008). Alternately, it is entirely possible that these structures are indeed ancestral but have undergone massive tissue reorganizations to reposition their attachment points. Such repositioning could be caused by moving the location of a critical signal or transcription factor within the tissues. Alternately, these structures could be specified before the discernable tissues of the phallus are separated, and their migration could be caused by differences in tissue folding. Under this scenario, we would anticipate that critical tissue patterning regulators of these processes are activated before these tissues become discernable. Finally, it is entirely possible that completely different networks account for the appearance of these unique structures. Developmental genetic analysis of the genes that produce the phallic processes described above will aid us in distinguishing these models. Recent work has identified several genes that are spatially restricted to the pregonites and postgonal processes of *D. melanogaster* providing an ideal set of candidates to examine (Vincent et al., 2019). Thus, we envision that detailed mechanisms of parallelism, repositioning, and novelty will emerge from studying systems where both network architecture is accessible, and genetic manipulations can be introduced to test the sufficiency of these mechanisms to produce these novel morphological structures.

## Acknowledgments

We would like to thank Deepak Dharmadhikari for his help with imaging, the Cornell Species and Bloomington Stock Centers for providing fly strains used in this study, Ben Vincent and the Rebeiz lab for their comments on the project and manuscript, Virginie Courtier-Orgogozo, Masanori Toda, Yoshitaka Kamimura and the Terminalia consortium for their insights on this project. We would also like to thank TaxoDros and the Japan Drosophila Database for their work cataloging resources for the original species descriptions for those analyzed in this study. This work was supported by a grant from the National Institutes of Health (R35GM141967 to M.R.). We want to give our heartfelt condolences to the family, friends, and colleagues of Jean David who passed away during publication of this work (Yassin, Gibert, & Capy, 2021). Jean was a truly inspirational scientist and friend; we were privileged to know him.

## Data Availability Statement

The data that support the findings of this study are available from the corresponding author upon reasonable request.

## Supplemental figures

**Figure S1:**
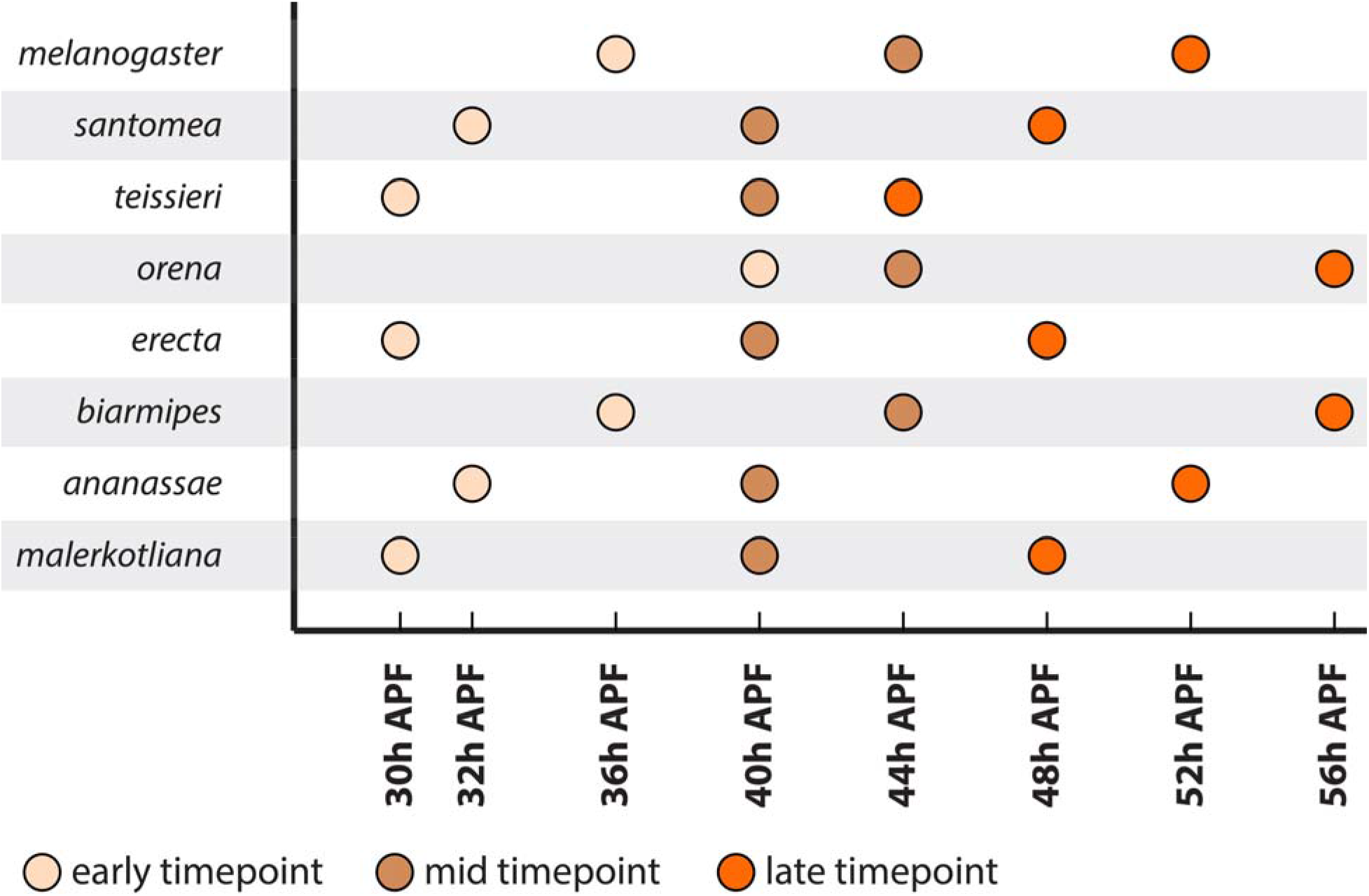
A summary of our designations of early (beige) mid (brown) and late (orange) developmental timepoints for each species. The early timepoint was designated when a cleavage between the epandrial ventral lobe (lateral plate) and surstylus (clasper) first forms. The mid timepoint was designated by when the cleavage of the epandrial ventral lobe and surstylus reached half of its total length. The late time point was designated as directly preceding when the cerci (anal plates) close over the gap between them.

**Figure S2:**
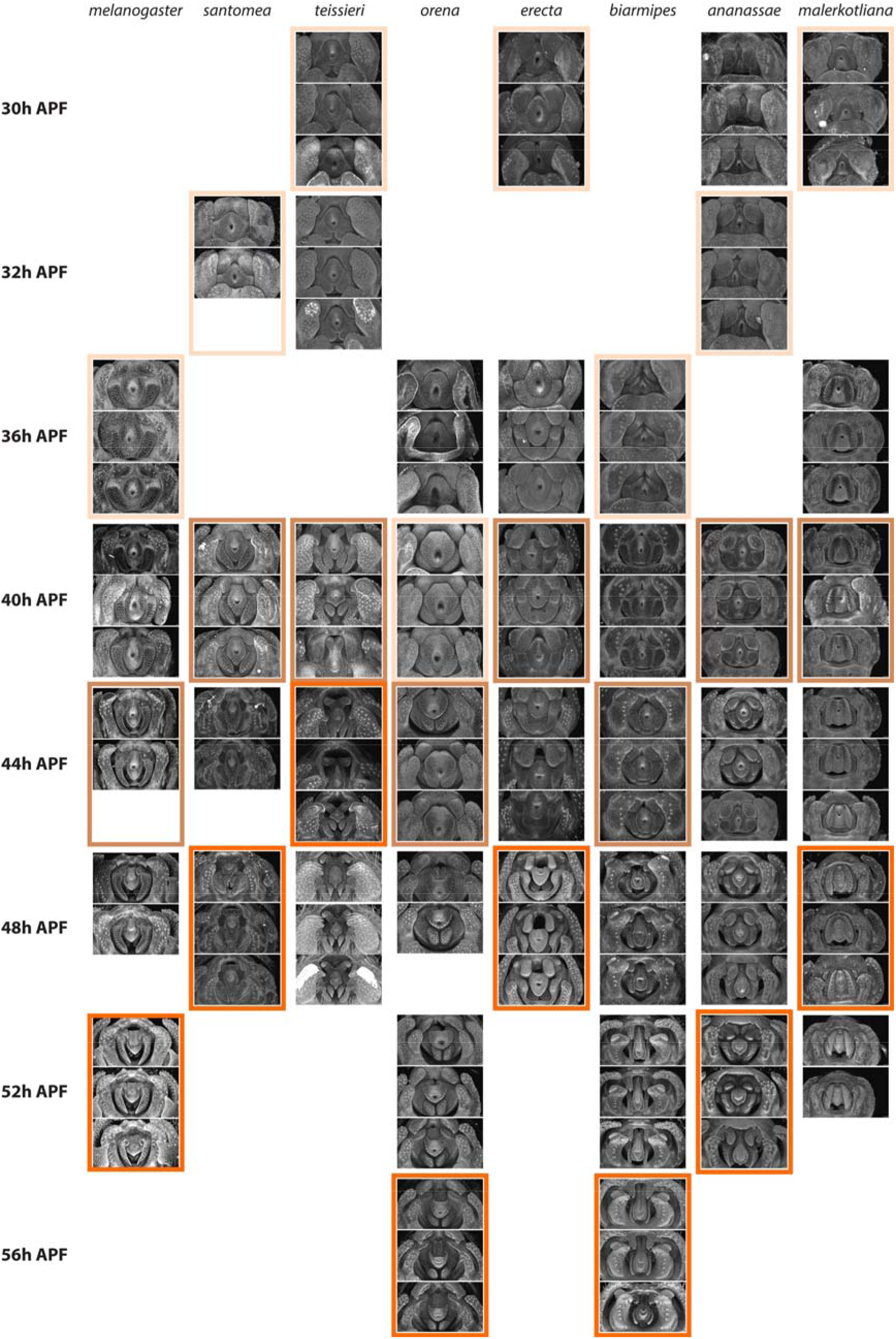
Full ECAD time course of ventral genitalia across the *melanogaster* species group. Developing pupal genitalia of *D. santomea, D. teissieri, D. orena, D. erecta, D. biarmipes, D. ananassae*, and *D. malerkotliana* stained for ECAD, apical cellular junctions, highlighting the overall morphology. Note that *D. melanogaster* samples use a transgenic line *arm-GFP* that also labels the apical cell junctions. Colored boxes highlight our three designated developmental timepoints for each species: early (beige) mid (brown) and late (orange) as shown in Figure S1.

**Figure S3:**
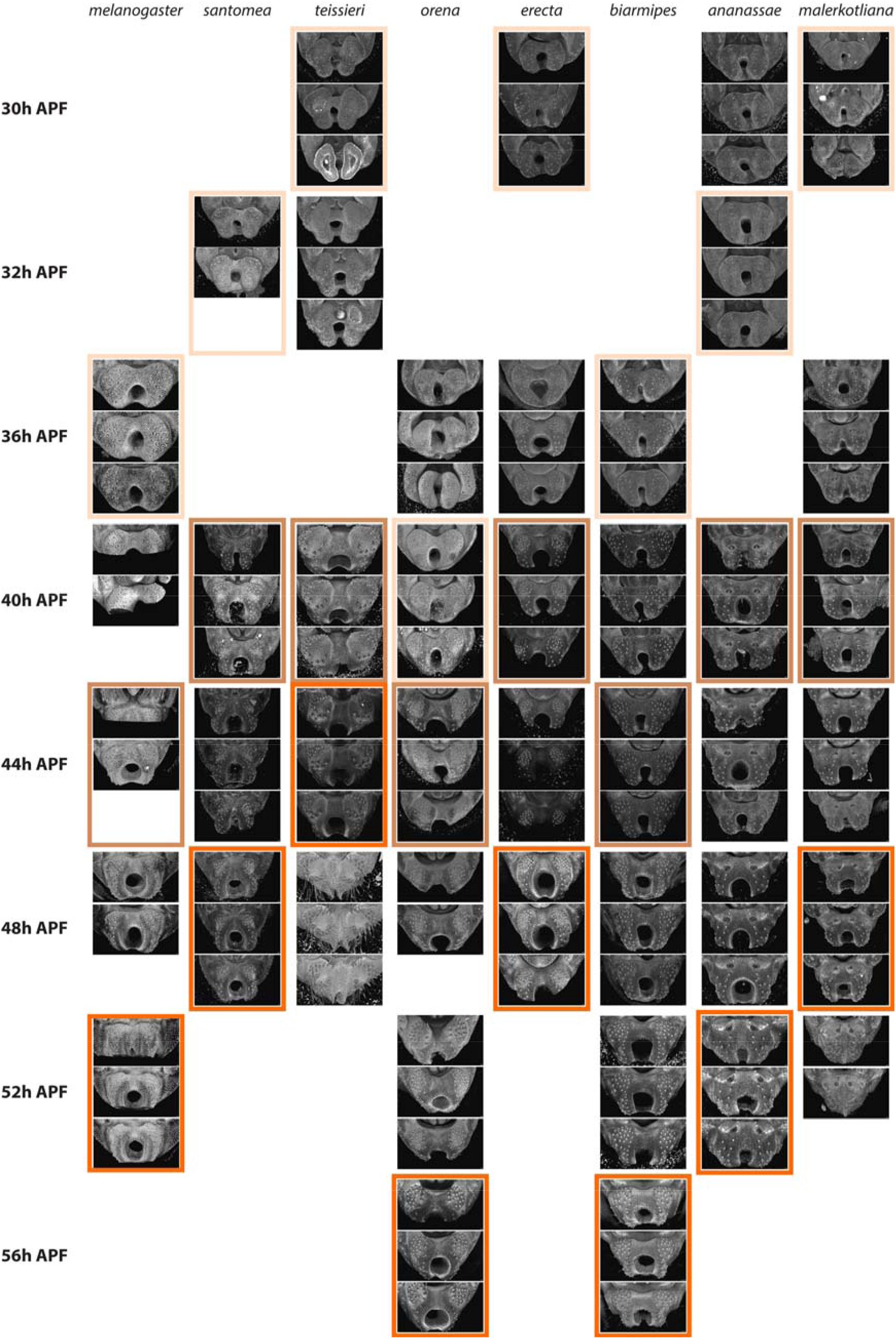
Full ECAD time course of dorsal genitalia and analia across *melanogaster* species group. Developing pupal genitalia of *D. santomea, D. teissieri, D. orena, D. erecta, D. biarmipes, D. ananassae*, and *D. malerkotliana* stained for ECAD, apical cellular junctions, highlighting the overall morphology. Note that *D. melanogaster* samples use a transgenic line arm-GFP that also labels the apical cell junctions. Colored boxes highlight our three designated developmental timepoints for each species: early (beige) mid (brown) and late (orange) as shown in Figure S1.

**Figure S4:**
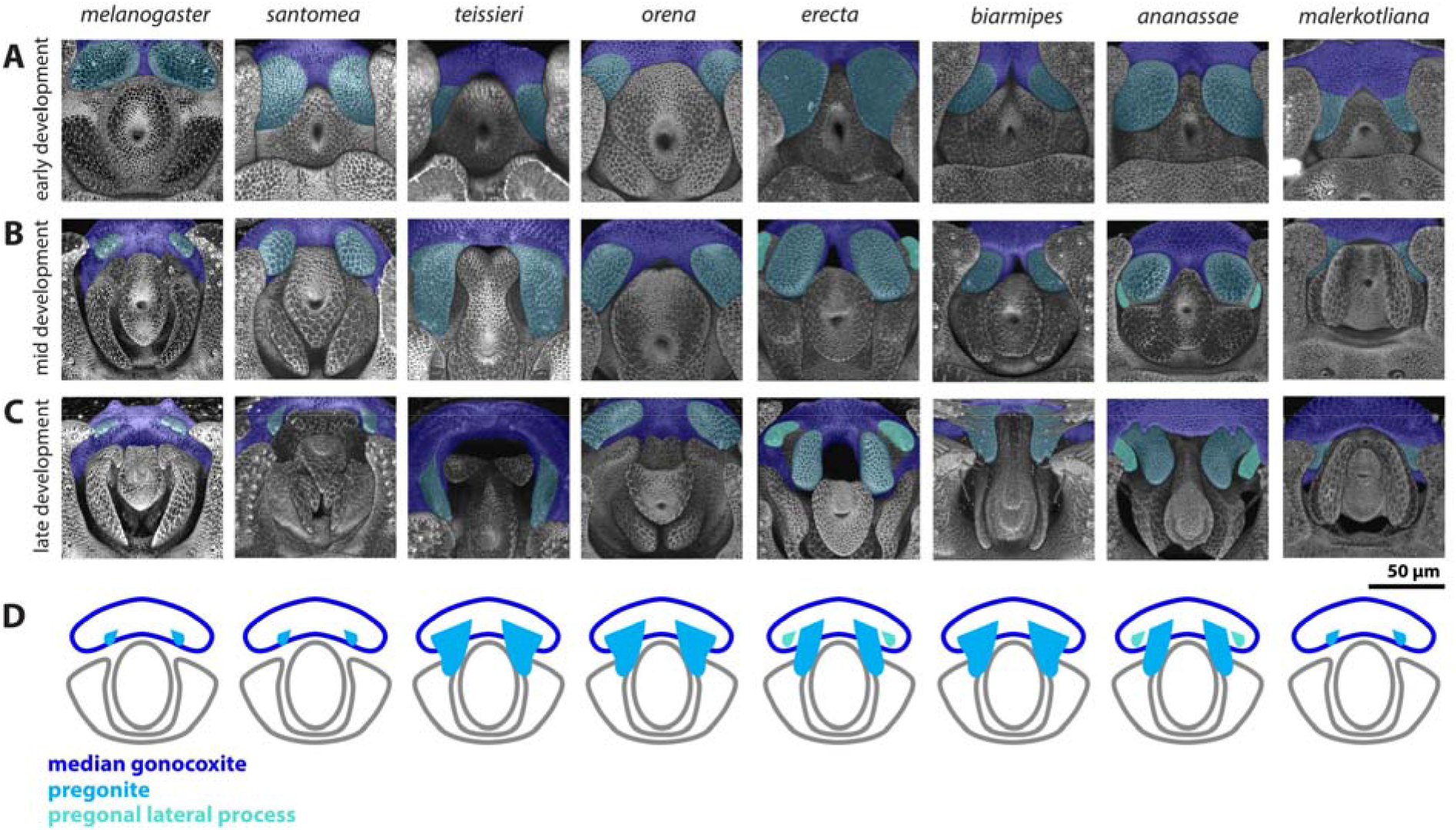
Developing ventral primordia of all analyzed species. **A-C)** The median gonocoxite is highlighted in dark blue, the pregonite is highlighted in with light blue, and the pregonal lateral process is highlighted in teal **D)** Schematic representation of the median gonocoxite (dark blue), the pregonite (light blue), pregonal lateral process (teal). Note that *D. melanogaster* samples use a transgenic line *arm-GFP*, while all other samples are stained for E-cadherin, both of which label apical cell junctions.

**Figure S5:**
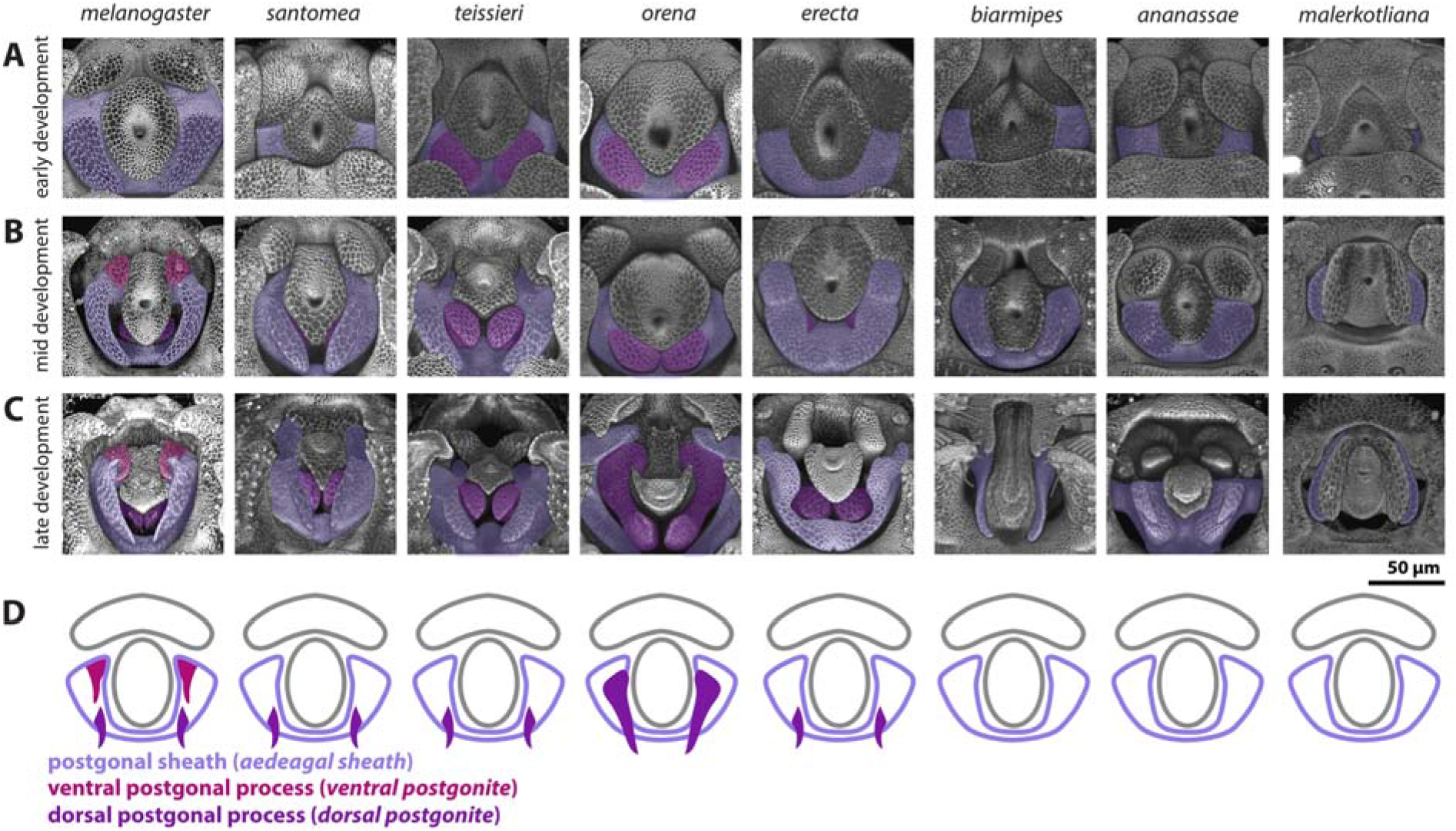
Developing dorsolateral primordia of all analyzed species. **A-C)** The postgonal sheath is highlighted in light purple, the ventral postgonal process is highlighted in magenta, and the dorsal postgonal process is highlighted in violet **D)** Schematic representations of the postgonal sheath (light purple), the ventral postgonal process (magenta), and dorsal postgonal process (violet). Aedeagal sheath is an alternative term for postgonal sheath, ventral postgonite is an alternative term for ventral postgonal process, and dorsal postgonite is an alternative term for dorsal postgonal process. Note that *D. melanogaster* samples use a transgenic line arm-GFP, while all other samples are stained for E-cadherin, both of which label apical cell junctions.

**Figure S6:**
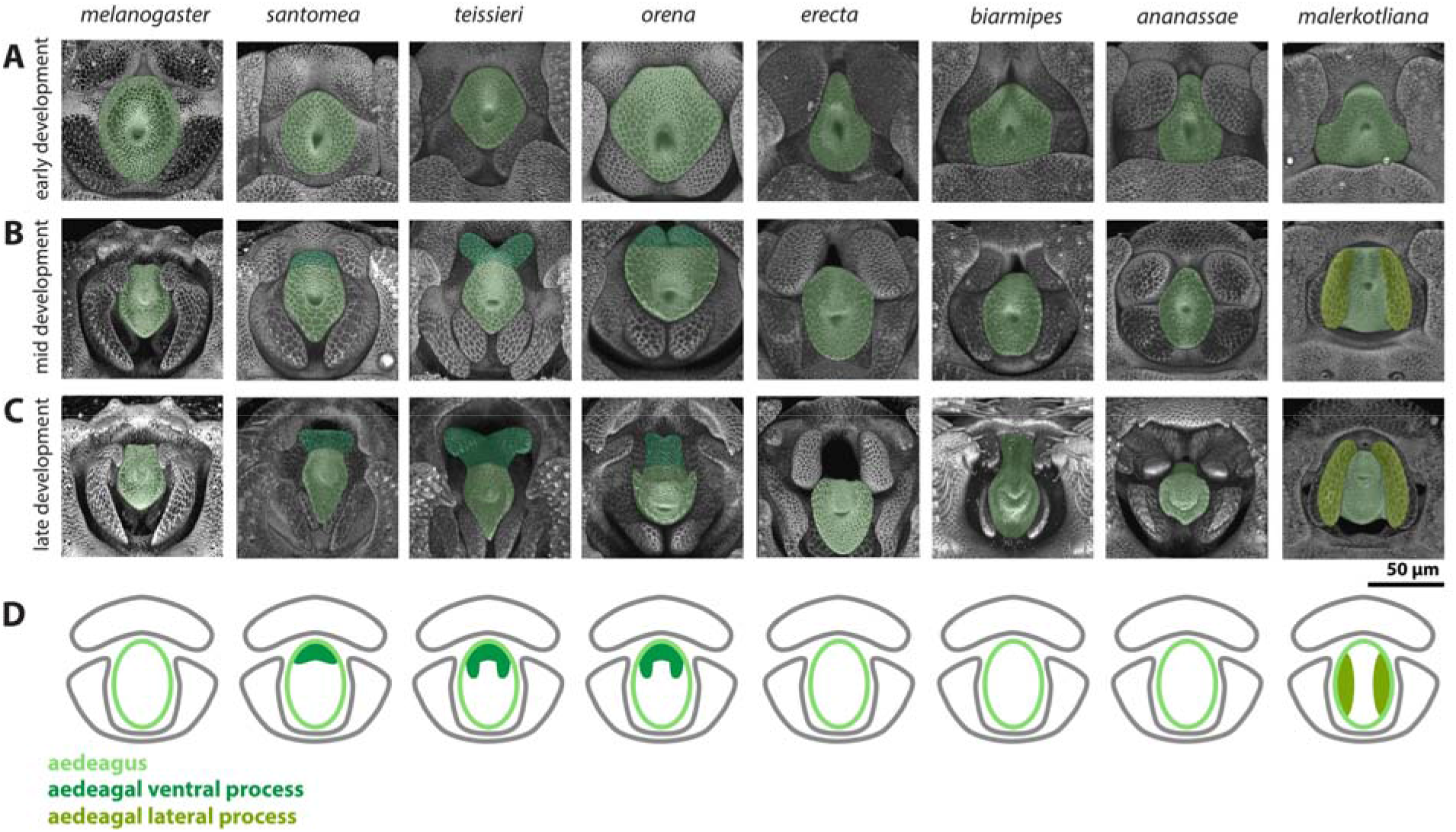
Developing central primordia of all analyzed species. **A-C)** The aedeagus is highlighted in light green, the aedeagal ventral process is highlighted in with dark green and the aedeagal lateral process is highlighted in yellow-green. **D)** Cartoon representations of the aedeagus (light green) aedeagal ventral process (dark green) and aedeagal lateral process (yellow-green). Note that *D. melanogaster* samples use a transgenic line *arm-GFP*, while all other samples are stained for E-cadherin, both of which label apical cell junctions.

**Figure S7:**
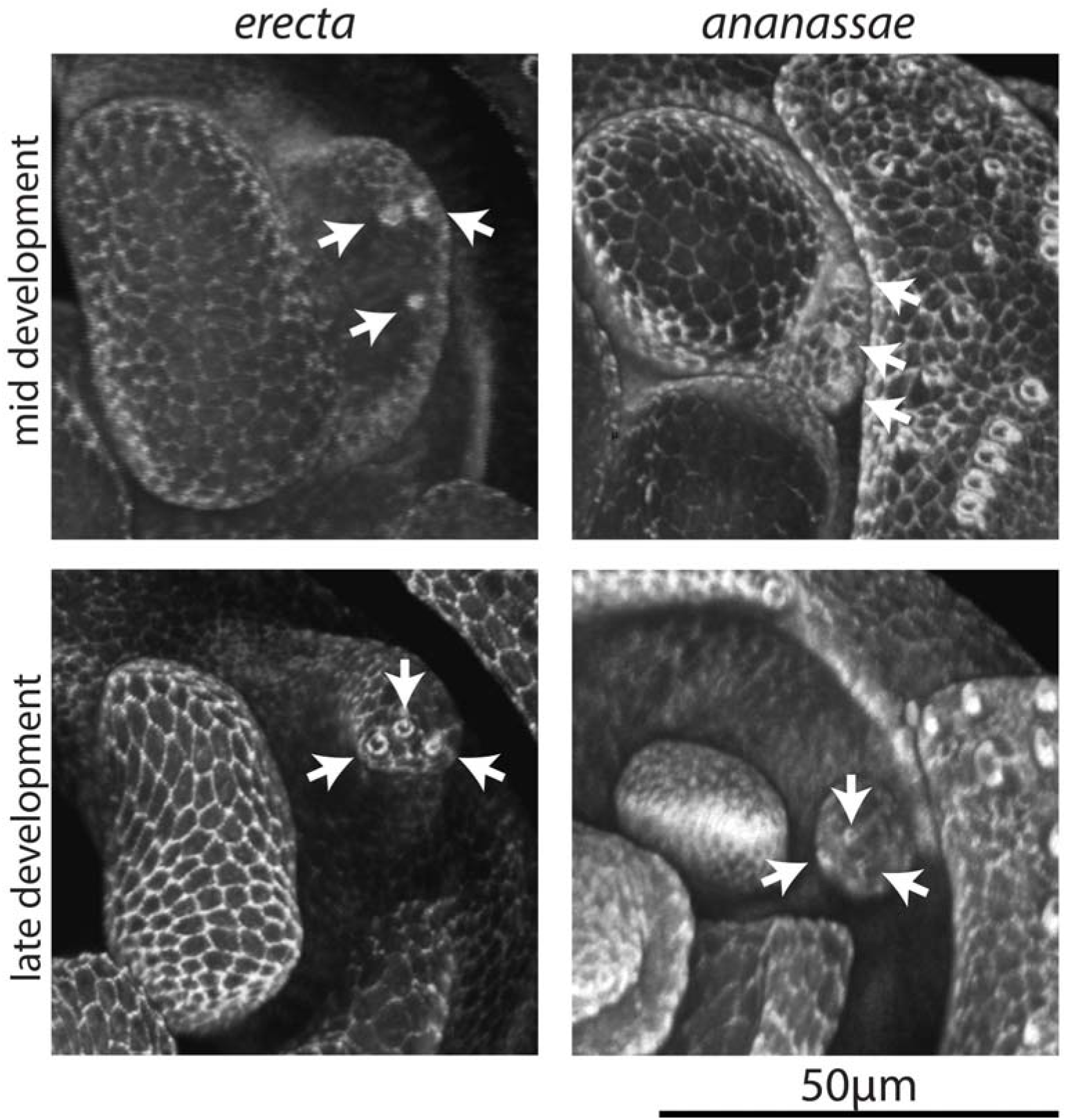
A zoom in on the developing pregonites of *D. erecta* and *D. ananassae*. Apical cell junctions are labeled through ECAD staining. White arrows indicate the location of the three developing pregonal bristles found on the pregonal lateral process.

## References

Abouheif, E. (2008). Parallelism as the pattern and process of mesoevolution. Evolution and Development, 10(1), 3–5. https://doi.org/10.1111/j.1525-142X.2007.00208.x

Acebes, A., Cobb, M., & Ferveur, J. F. (2003). Species-specific effects of single sensillum ablation on mating position in Drosophila. Journal of Experimental Biology, 206(17), 3095–3100. https://doi.org/10.1242/jeb.00522

Ah-King, M., Barron, A. B., & Herberstein, M. E. (2014). Genital Evolution: Why Are Females Still Understudied? PLoS Biology, 12(5), 1–7. https://doi.org/10.1371/journal.pbio.1001851

Ahmad, S. M., & Baker, B. S. (2002). Sex-Specific Deployment of FGF Signaling in Drosophila Recruits Mesodermal Cells into the Male Genital Imaginal Disc, 109, 651–661.

Al Sayad, S., & Yassin, A. (2019). Quantifying the extent of morphological homoplasy: A phylogenetic analysis of 490 characters in Drosophila. Evolution Letters, 3(3), 286–298. https://doi.org/10.1002/evl3.115

Aspiras, A. C., Smith, F. W., & Angelini, D. R. (2011). Sex-specific gene interactions in the patterning of insect genitalia. Developmental Biology, 360(2), 369–380. https://doi.org/10.1016/j.ydbio.2011.09.026

Avila, F. W., Sirot, L. K., Laflamme, B. A., Rubinstein, C. D., & Wolfner, M. F. (2011). Insect seminal fluid proteins: Identification and function. Annual Review of Entomology, 56, 21–40. https://doi.org/10.1146/annurev-ento-120709-144823

Bächli, G., Vilela, C., Andersson Escher, S., & Saura, A. (2004). The Drosophilidae (Diptera) of 252 Fennoscandia and Denmark. Fauna Entomologica Scandinavica. Brill.

Bock, I. R., & Wheeler, M. R. (1972). The Drosophila melanogaster species group. The University of Texas Publication, VII(7213), 1–102.

Brennan, P. L. R., & Prum, R. O. (2015). Mechanisms and evidence of genital coevolution: The roles of natural selection, mate choice, and sexual conflict. Cold Spring Harbor Perspectives in Biology, 7(7), 1–21. https://doi.org/10.1101/cshperspect.a017749

Bruce, H. S., & Patel, N. H. (2020). Knockout of crustacean leg patterning genes suggests that insect wings and body walls evolved from ancient leg segments. Nature Ecology and Evolution, 4(12), 1703–1712. https://doi.org/10.1038/s41559-020-01349-0

Bryant, P. J., & Hsei, B. W. (1977). Pattern formation in asymmetrical and symmetrical imaginal discs of Drosophila melanogaster. American Zoologist, 17(3), 595–611. https://doi.org/10.1093/icb/17.3.595

Church, S. H., & Extavour, C. G. (2020). Null hypotheses for developmental evolution. Development (Cambridge), 147(8), 1–6. https://doi.org/10.1242/DEV.178004

Clark-Hachtel, C. M., & Tomoyasu, Y. (2020). Two sets of candidate crustacean wing homologues and their implication for the origin of insect wings. Nature Ecology and Evolution, 4(12), 1694–1702. https://doi.org/10.1038/s41559-020-1257-8

Cordero-Rivera, A. (2017). Sexual conflict and the evolution of genitalia: male damselflies remove more sperm when mating with a heterospecific female. Scientific Reports, 7(1), 1–8. https://doi.org/10.1038/s41598-017-08390-3

de Reuille, P. B., Routier-Kierzkowska, A. L., Kierzkowski, D., Bassel, G. W., Schüpbach, T., Tauriello, G.,… Smith, R. S. (2015). MorphoGraphX: A platform for quantifying morphogenesis in 4D. ELife, 4(MAY). https://doi.org/10.7554/eLife.05864

Doleschall, C. L. (1858). Derde bijdrage tot de kennis der Dipteren fauna van Nederlandsch Indie. Natuurkundig Tijdschrift Voor Nederlandsch Indie, 17, 73–128.

Eberhard, W. G. (1985). Sexual Selection and Animal Genitalia. Cambridge, Mass: Harvard University Press.

Emlen, D. J., Szafran, Q., Corley, L. S., & Dworkin, I. (2006). Insulin signaling and limb-patterning: Candidate pathways for the origin and evolutionary diversification of beetle “horns.” Heredity, 97(3), 179–191. https://doi.org/10.1038/sj.hdy.6800868

Epper, F. (1983). The Evagination of the Genital Imaginal Discs of Drosophila melanogaster II. Morphogenesis of the Intersexual Genital Disc of the Mutant doublesex-dominant (dsx D). Roux’s Archives of Developmental Biology, 192, 280–284.

Glassford, W. J., Johnson, W. C., Dall, N. R., Smith, S. J., Liu, Y., Boll, W.,… Rebeiz, M. (2015). Co-option of an Ancestral Hox-Regulated Network Underlies a Recently Evolved Morphological Novelty Article. Developmental Cell, 34(5), 520–531. https://doi.org/10.10l6/j.devcel.2015.08.005

Green, J. E., Cavey, M., Médina Caturegli, E., Aigouy, B., Gompel, N., & Prud’homme, B. (2019). Evolution of Ovipositor Length in Drosophila suzukii Is Driven by Enhanced Cell Size Expansion and Anisotropic Tissue Reorganization. Current Biology, 29(12), 2075–2082.e6. https://doi.org/10.1016/j.cub.2019.05.020

Hagen, J. F. D., Mendes, C. C., Blogg, A., Payne, A., Tanaka, K. M., Gaspar, P.,… Nunes, M. D. S. (2019). Tartan underlies the evolution of Drosophila male genital morphology. Proceedings of the National Academy of Sciences of the United States of America, 116(38), 19025–19030. https://doi.org/10.1073/pnas.1909829116

Hagen, J. F. D., Mendes, C. C., Booth, S. R., Figueras Jimenez, J., Tanaka, K. M., Franke, F. A.,… McGregor, A. P. (2021). Unraveling the Genetic Basis for the Rapid Diversification of Male Genitalia between Drosophila Species. Molecular Biology and Evolution, 38(2), 437–448. https://doi.org/10.1093/molbev/msaa232

Hinman, V. F., Nguyen, A. T., Cameron, R. A., & Davidson, E. H. (2003). Developmental gene regulatory network architecture across 500 million years of echinoderm evolution. Proceedings of the National Academy of Sciences of the United States of America, 100(23), 13356–13361. https://doi.org/10.1073/pnas.2235868100

Hosken, D. J., Archer, C. R., House, C. M., & Wedell, N. (2019). Penis evolution across species: divergence and diversity. Nature Reviews Urology, 16(2), 98–106. https://doi.org/10.1038/s41585-018-0112-z

Hsu, T. C. (1949). The external genital apparatus of male Drosophilidae in relation to systematics. The University of Texas Publication, 4920, 80–142.

Huang, J., Huang, L., Chen, Y. J., Austin, E., Devor, C. E., Roegiers, F., & Hong, Y. (2012). Differential regulation of adherens junction dynamics during apical-basal polarization. Development, 139(3), 4001–4013. https://doi.org/10.1242/jcs.086694

Huber, B. A. (2005). Sexual selection research on spiders: Progress and biases. Biological Reviews of the Cambridge Philosophical Society, 80(3), 363–385. https://doi.org/10.1017/S1464793104006700

Jagadeeshan, S., & Singh, R. S. (2006). A time-sequence functional analysis of mating behaviour and genital coupling in Drosophila: Role of cryptic female choice and male sex-drive in the evolution of male genitalia. Journal of Evolutionary Biology, 19(4), 1058–1070. https://doi.org/10.1111/j.1420-9101.2006.01099.x

Kamimura, Y. (2007). Twin intromittent organs of Drosophila for traumatic insemination. Biology Letters, 3(4), 401–404. https://doi.org/10.1098/rsbl.2007.0192

Kamimura, Y. (2010). Copulation anatomy of Drosophila melanogaster (Diptera: Drosophilidae): Wound-making organs and their possible roles. Zoomorphology, 129(3), 163–174. https://doi.org/10.1007/s00435-010-0109-5

Kamimura, Y. (2012). Correlated evolutionary changes in Drosophila female genitalia reduce the possible infection risk caused by male copulatory wounding. Behavioral Ecology and Sociobiology, 66(8), 1107–1114. https://doi.org/10.1007/s00265-012-1361-0

Kamimura, Y. (2016). Significance of constraints on genital coevolution: Why do female Drosophila appear to cooperate with males by accepting harmful matings? Evolution; International Journal of Organic Evolution, 70(7), 1674–1683. https://doi.org/10.1111/evo.12955

Kamimura, Y., & Mitsumoto, H. (2011). Comparative copulation anatomy of the Drosophila melanogaster species complex (Diptera: Drosophilidae). Entomological Science, 14(4), 399–410. https://doi.org/10.1111/j.1479-8298.2011.00467.x

Kamimura, Y., & Mitsumoto, H. (2012a). Genital coupling and copulatory wounding in drosophila teissieri (diptera: Drosophilidae). Canadian Journal of Zoology, 90(12), 1437–1440. https://doi.org/10.1139/cjz-2012-0186

Kamimura, Y., & Mitsumoto, H. (2012b). Lock-and-key structural isolation between sibling Drosophila species. Entomological Science, 15(2), 197–201. https://doi.org/10.1111/j.1479-8298.2011.00490.x

Kamimura, Y., & Polak, M. (2011). Does surgical manipulation of drosophila intromittent organs affect insemination success? Proceedings of the Royal Society B: Biological Sciences, 278(1707), 815–816. https://doi.org/10.1098/rspb.2010.2431

Klaus, A. V., Kulasekera, V. L., & Schawaroch, V. (2003). Three-dimensional visualization of insect morphology using confocal laser scanning microscopy. Journal of Microscopy, 212(2), 107–121. https://doi.Org/10.1046/j.1365-2818.2003.01235.x

Lachaise, D., Capy, P., Cariou, M.-L., Joly, D., Lemeunier, F., & David, J. R. (2004). Nine relatives from one African ancestor: population biology and evolution of the Drosophila melanogaster subgroup species. In R. S. Singh, M. K. Uyenoyama, & S. K. Jain (Eds.), The Evolution of Population Biology (pp. 315–344). Cambridge University Press. https://doi.org/10.1017/cbo9780511542619.019

Lachaise, D., Harry, M., Solignac, M., Lemeunier, F., Benassi, V., & Cariou, M. L. (2000). Evolutionary novelties in islands: Drosophila santomea, a new melanogaster sister species from Sao Tome. Proceedings of the Royal Society B: Biological Sciences, 267(1452), 1487–1495. https://doi.org/10.1098/rspb.2000.1169

Linz, D. M., Hu, Y., & Moczek, A. P. (2020). From descent with modification to the origins of novelty. Zoology, 143(August), 125836. https://doi.org/10.10l6/j.zool.2020.125836

Malloch, J. R. (1924). Two Drosophilidae from Coimbatore. Memoirs of the Department of Agriculture in India. Entomological Series, 8(6), 63–65.

Markow, T. A., & O’Grady, P. M. (2006). Drosophila: A Guide to Species Identification and Use. Academic Press. London.

Masly, J. P. (2012). 170 Years of “Lock-and-Key”: Genital Morphology and Reproductive Isolation. International Journal of Evolutionary Biology, 2012, 1–10. https://doi.org/10.1155/2012/247352

Masly, J. P., & Kamimura, Y. (2014). Asymmetric mismatch in strain-specific genital morphology causes increased harm to drosophila females. Evolution, 68(8), 2401–2411. https://doi.org/10.1111/evo.12436

Mattei, A. L., Riccio, M. L., Avilaa, F. W., Wolfner, M. F., & Denlinger, D. L. (2015). Integrated 3D view of postmating responses by the Drosophila melanogaster female reproductive tract, obtained by micro-computed tomography scanning. Proceedings of the National Academy of Sciences of the United States of America, 112(27), 8475–8480. https://doi.org/10.1073/pnas.1505797112

McNamara, K. B., Dougherty, L. R., Wedell, N., & Simmons, L. W. (2019). Experimental evolution reveals divergence in female genital teeth morphology in response to sexual conflict intensity in a moth. Journal of Evolutionary Biology, 32(5), 519–524. https://doi.org/10.1111/jeb.13428

McPeek, M. A., Shen, L., & Farid, H. (2009). The correlated evolution of three-dimensional reproductive structures between male and female damselflies. Evolution, 63(1), 73–83. https://doi.org/10.1111/j.1558-5646.2008.00527.x

Meigen, J. W. (1830). Systematische Beschreibung der bekannten europäischen zweiflugeligen Insekten. Schulze.

Moczek, A. P. (2008). On the origins of novelty in development and evolution. BioEssays, 30(5), 432–447. https://doi.org/10.1002/bies.20754

Muto, L., Kamimura, Y., Tanaka, K. M., & Takahashi, A. (2018). An innovative ovipositor for niche exploitation impacts genital coevolution between sexes in a fruit-damaging Drosophila. Proceedings of the Royal Society B: Biological Sciences. https://doi.org/10.1098/rspb.2018.1635

Nagy, O., Nuez, I., Savisaar, R., Peluffo, A. E., Yassin, A., Lang, M.,… Courtier-Orgogozo, V. (2018). Correlated Evolution of Two Copulatory Organs via a Single cis-Regulatory Nucleotide Change. Current Biology, 28(21), 1–8. https://doi.org/10.1016/j.cub.2018.08.047

Obbard, D. J., MacLennan, J., Kim, K. W., Rambaut, A., O’Grady, P. M., & Jiggins, F. M. (2012). Estimating divergence dates and substitution rates in the drosophila phylogeny. Molecular Biology and Evolution, 29(11), 3459–3473. https://doi.org/10.1093/molbev/mssl50

Okada, T. (1954). Comparative morphology of the drosophilid flies. I. Phallic organs of the melanogaster group. Kontyu, 22, 36–46.

Parshad, R., & Paika, I. J. (1964). Drosophilid survey of India. II. Taxonomy and cytology of the subgenus Sophophora (Drosophila). Research Bulletin of the Panjab University. Science., 15, 225–252.

Peluffo, A. E., Nuez, I., Debat, V., Savisaar, R., Stern, D. L., & Orgogozo, V. (2015). A major locus controls a genital shape difference involved in reproductive isolation between Drosophila yakuba and Drosophila santomea. G3: Genes, Genomes, Genetics, 5(12), 2893–2901. https://doi.org/10.1534/g3.115.023481

Peluffo, A., Hamdani, M., Vargas-Valderrama, A., David, J., Mallard, F., Graner, F., & Courtier-Orgogozo, V. (2021). A morphological trait involved in reproductive isolation between Drosophila sister species is sensitive to temperature. BioRx. https://doi.org/10.1101/2020.01.20.911826

Perry, J. C., & Rowe, L. (2018). Sexual conflict in its ecological setting. Philosophical Transactions of the Royal Society B: Biological Sciences, 373(1757). https://doi.org/10.1098/rstb.2017.0418

Prud’Homme, B., Minervino, C., Hocine, M., Cande, J. D., Aouane, A., Dufour, H. D.,… Gompel, N. (2011). Body plan innovation in treehoppers through the evolution of an extra wing-like appendage. Nature, 473(7345), 83–86. https://doi.org/10.1038/nature09977

Quade, F. S. C., Holtzheimer, J., Frohn, J., Töpperwien, M., Salditt, T., & Prpic, N. M. (2019). Formation and development of the male copulatory organ in the spider Parasteatoda tepidariorum involves a metamorphosis-like process. Scientific Reports, 9(1), 1–12. https://doi.org/10.1038/s41598-019-43192-9

Rebeiz, M., Patel, N. H., & Hinman, V. F. (2015). Unraveling the Tangled Skein: The Evolution of Transcriptional Regulatory Networks in Development. Annual Review of Genomics and Human Genetics, 16(1), 103–131. https://doi.org/10.1146/annurev-genom-091212-153423

Rebeiz, M., & Tsiantis, M. (2017). Enhancer evolution and the origins of morphological novelty. Current Opinion in Genetics and Development, 45, 115–123. https://doi.org/10.1016/j.gde.2017.04.006

Rhebergen, F. T., Courtier-Orgogozo, V., Dumont, J., Schilthuizen, M., & Lang, M. (2016). Drosophila pachea asymmetric lobes are part of a grasping device and stabilize one-sided mating. BMC Evolutionary Biology, 16(1), 1–24. https://doi.org/10.1186/sl2862-016-0747-4

Rice, G., David, J. R., Kamimura, Y., Masly, J. P., Alistair, P., Nagy, O.,… Yassin, A. (2019). A standardized nomenclature and atlas of the male terminalia of Drosophila melanogaster. Fly, 13(1-4), 51–64. https://doi.org/10.1080/19336934.2019.1653733

Robertson, H. (1988). Mating Asymmetries and Phylogeny in the Drosophila melanogaster Species Complex, 42.

Rowe, L., & Arnqvist, G. (2012). Sexual selection and the evolution of genital shape and complexity in water striders. Evolution, 66(1), 40–54. https://doi.org/10.1111/j.1558-5646.2011.01411.x

Sánchez, V., & Cordero, C. (2014). Sexual coevolution of spermatophore envelopes and female genital traits in butterflies: Evidence of male coercion? PeerJ, 2014(1), 1–12. https://doi.org/10.7717/peerj.247

Shao, L., Chung, P., Wong, A., Siwanowicz, I., Kent, C. F., Long, X., & Heberlein, U. (2019). A Neural Circuit Encoding the Experience of Copulation in Female Drosophila. Neuron, 102(5), 1025–1036.E6. https://doi.Org/10.1016/j.neuron.2019.04.009

Simmons, L. W. (2014). Sexual selection and genital evolution. Austral Entomology, 53(1), 1–17. https://doi.org/10.1111/aen.12053

Simmons, L. W., & Fitzpatrick, J. L. (2019). Female genitalia can evolve more rapidly and divergently than male genitalia. Nature Communications, 10(1), 1–8. https://doi.org/10.1038/s41467-019-09353-0

Smith, S. J., Davidson, L. A., & Rebeiz, M. (2020). Evolutionary expansion of apical extracellular matrix is required for the elongation of cells in a novel structure. ELife, 9. https://doi.org/10.7554/eLife.55965

Steedman, H. F. (1958). Dimethyl Hydantoin Formaldehyde: A new Water-soluble Resin for Use as a Mounting Medium. Journal of Cell Science, 99(4), 451–452. https://doi.org/10.1163/187529266X00220

Tanaka, K., Barmina, O., & Kopp, A. (2009). Distinct developmental mechanisms underlie the evolutionary diversification of Drosophila sex combs. Proceedings of the National Academy of Sciences, 106(12), 4764–4769. https://doi.org/10.1073/pnas.0807875106

Tsacas, L. (1971). Drosophila teissieri, nouvelle espece africaine du groupe melanogaster et note sur deux autres especes nouvelles pour I’Afrique (Dipt. Drosophilidae). Bulletin de La Société Entomologigue de France, 76, 35–45.

Tsacas, L., Bocquet, C., Daguzan, M., & Mercier, A. (1971). Comparaison des genitalia males de Drosophila melanogaster, de Drosophila simulans et de leurs hybrids. Annales de La Société Entomologigue de France, 7, 75–93. Retrieved from https://ci.nii.ac.jp/naid/10030580910/

Tsacas, L., & David, J. (1978). Une septieme espece appartenant au sous-groupe Drosophila melanogaster Meigen: Drosophila orena spec. nov. du Cameroun. (Diptera: Drosophilidae). Beiträge Zur Entomologie, 28, 179–182.

Tsacas, L., & Lachaise, D. (1974). Quatre nouvelles especes de la Cote-d?voire du genre Drosophila, groupe melanogaster, et discussion de I’origine du sous-groupe melanogaster (Diptera: Drosophilidae). Annales de I’Université d’Abidjan Série E: Ecologie, 7, 193–211.

van Emden, F., & Hennig, W. (1970). Taxonomists’ glossary of genitalia of insects. (S. L. Tuxen, Ed.) (2nd ed.). Munksgaard, Copenhagen.

Vincent, B. J., Rice, G. R., Wong, G. M., Glassford, W. J., Downs, K. I., Shastay, J. L.,… Rebeiz, M. (2019). An atlas of transcription factors expressed in the Drosophila melanogaster pupal terminalia. G3: Genes/Genomes/Genetics, 9(December), 3961–3972. https://doi.org/10.1101/677260

Wagner, G. (2007). The developmental genetics of homology. Nature Reviews Genetics, 8(6), 473–479. https://doi.org/10.1038/nrg2099

Wagner, G. P., & Lynch, V. J. (2010). Evolutionary novelties. Current Biology, 20(2), 48–52. https://doi.org/10.10l6/j.cub.2009.11.010

Yassin, A., Gibert, P., & Capy, P. (2021). In memoriam: Jean R. David (1931-2021). Retrieved from https://genestogenomes.org/in-memoriam-jean-r-david-1931-2021/

Yassin, A., & Orgogozo, V. (2013). Coevolution between Male and Female Genitalia in the Drosophila melanogaster Species Subgroup. PLoS ONE, 8(2). https://doi.org/10.1371/journal.pone.0057158

